# A potential space making role for a lytic transglycosylase in cell wall biogenesis revealed by a beta-lactamase induction phenotype in *Pseudomonas aeruginosa*

**DOI:** 10.1101/2023.09.05.556436

**Authors:** Coralie Fumeaux, Thomas G. Bernhardt

**Affiliations:** Department of Microbiology, Harvard Medical School, Boston, MA, USA; Institute of Microbiology, Lausanne University Hospital, University of Lausanne, 1011 Lausanne, Switzerland; Howard Hughes Medical Institute, Chevy Chase, MD 20815, USA

## Abstract

*Pseudomonas aeruginosa* encodes the beta-lactamase AmpC, which promotes resistance to beta-lactam antibiotics. Expression of *ampC* is induced by fragments of the peptidoglycan (PG) cell wall released upon beta-lactam treatment. These drugs target transpeptidase enzymes that form cell wall crosslinks. However, they do not block the activity of the transglycosylases that polymerize the glycan chains. Thus, drug-treated cells produce uncrosslinked PG polymers that have been shown to be rapidly degraded in the related gram-negative bacterium *Escherichia coli*. This degradation is performed by enzymes called lytic transglycosylases (LTs), which generate the anhydro-muropeptide (AMP) products sensed by the AmpR regulator that activates *ampC* expression. To identify factors required for proper PG biogenesis in *P. aeruginosa*, we used a reporter gene fusion to the *ampC* promoter to screen for mutants induced for *ampC* expression in the absence of drug. To our surprise, we found that inactivation of SltB1, an LT enzyme expected to produce the AMP products required for *ampC* induction, counterintuitively led to elevated *ampC* expression. This induction required another LT enzyme called MltG, suggesting that inactivation of SltB1 reduces the efficiency of PG crosslinking, causing the degradation of a subset of nascent PG strands by MltG to generate the inducing signal. Our results therefore support a model in which SltB1 uses its LT activity to open space in the PG matrix for the efficient insertion of new material, a function commonly thought to be restricted to endopeptidases that cut cell wall crosslinks.

**IMPORTANCE:** Inducible beta-lactamases like the *ampC* system of *Pseudomonas aeruginosa* are a common determinant of beta-lactam resistance among gram-negative bacteria. The regulation of *ampC* is elegantly tuned to detect defects in cell wall synthesis caused by beta-lactam drugs. Studies of mutations causing *ampC* induction in the absence of drug therefore promise to reveal new insights into the process of cell wall biogenesis in addition to aiding our understanding of how resistance to beta-lactam antibiotics arises in the clinic. In this study, an *ampC* induction phenotype for a mutant lacking an enzyme that cleaves cell wall glycans was used to uncover a potential role for glycan cleavage in making space in the wall matrix for the insertion of new material during cell growth.

## INTRODUCTION

*Pseudomonas aeruginosa* is an opportunistic pathogen that encodes multiple drug resistance mechanisms (1, 2). Infections with this bacterium can therefore be difficult to treat. One of its major resistance determinants is the AmpC beta-lactamase, which limits the effectiveness of many beta-lactam antibiotics against *P. aeruginosa* (3). Expression of the *ampC* gene is regulated in response to drug treatment. In the absence of antibiotic, it is expressed at low levels. However, treatment with some beta-lactams like cefoxitin, referred to as beta-lactamase inducers, results in potent induction of *ampC* expression and resistance. Other beta-lactams like piperacillin and ceftazidime are not *ampC* inducers (4). These drugs therefore have anti-pseudomonal activity despite the ability of AmpC to hydrolyze them. Mutants with defects in *ampC* regulation causing constitutive beta-lactamase production are resistant to piperacillin and ceftazidime. They are known to arise in the clinic and can result in treatment failures (5–9). There has thus been considerable interest in understanding *ampC* regulation and the mechanism by which mutations promote its aberrant overexpression.

The expression level of *ampC* is linked to the status of peptidoglycan (PG) synthesis and responds to signals produced when beta-lactam antibiotics disrupt the process (3, 10–13). The PG cell wall surrounds most bacteria and is essential for maintaining cellular integrity. It is composed of glycan chains with a repeating disaccharide unit of N-acetylmuramic acid (MurNAc) and N-acetylglucosamine (GlcNAc). A pentapeptide with the sequence L-Ala-ɣ-D-Glu-meso-diaminopimelic acid (mDAP)-D-Ala-D-Ala is attached to the MurNAc sugar. It is used to form crosslinks between glycan chains, generating the matrix-like structure of the wall (**Fig. 1**) (14). Two different types of synthases build the PG layer. The class A penicillin-binding proteins (aPBPs) possess both glycosyltransferase (GTase) and transpeptidase (TPase) activities in a single polypeptide for the polymerization and crosslinking of PG, respectively. The other major synthases are composed of complexes between a SEDS-family protein with GTase activity and a class B penicillin-binding protein (bPBP) with TPase activity (15–17). Because the PG matrix is continuous, the insertion of new material requires the action of PG cleaving enzymes to make space for the incoming nascent glycans (14). Currently, endopeptidases that cut the peptide crosslinks are the only known PG processing enzymes implicated as space-makers (14, 18–21).

**Figure 1.**
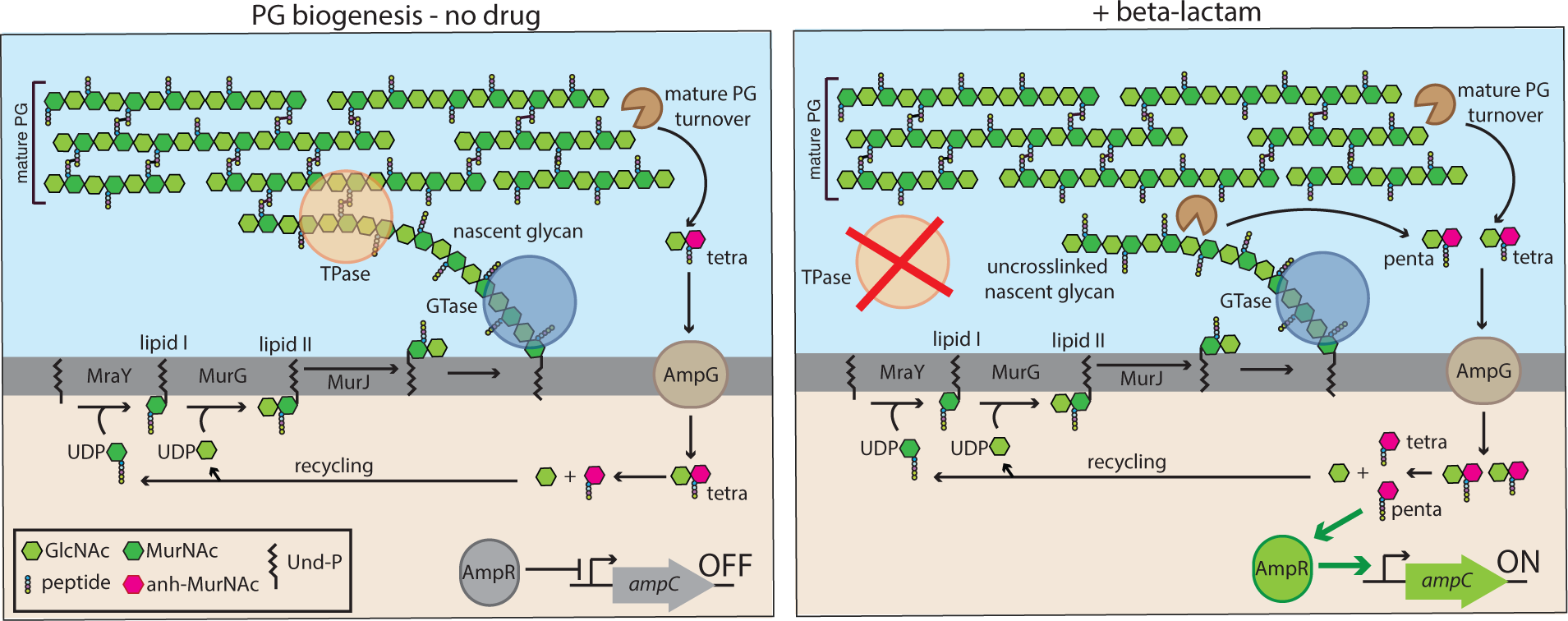
Overview of PG synthesis and *ampC* regulation. (Left) The PG matrix consists of glycan chains with the repeating unit of N-acetylmuramic acid (MurNAc) and N-acetylglucosamine (GlcNAc). Attached to the MurNAc sugars is a pentapeptide used to form crosslinks between adjacent glycans. PG synthesis starts in the cytoplasm, followed by the generation of lipid-linked precursors. Polymerization (GTase) and crosslinking (TPase) reactions at the membrane surface are used to form and insert nascent glycans into the mature matrix. The crosslinking reaction and carboxypeptidase enzymes act to rapidly convert the pentapeptide side chains to tetrapeptides via removal of the terminal D-Ala. The mature PG is subject to degradation by lytic transglycosylases (LT) and endopeptidases (EP) to generate anydro-MurNAc (anh-MurNAc) containing muropeptide (AMP) turnover products, which are imported to the cytoplasm by AmpG and recycled. The tetrapeptide AMP products are not thought to be good inducers for the activation of *ampC* expression by AmpR such that *ampC* is repressed in the absence of drug (48). (**Right**) Upon beta-lactam treatment, TPases are inhibited and uncrosslinked nascent glycans are formed. These glycans are rapidly degraded generating pentapeptide AMP products that are also transported to the cytoplasm by AmpG (36, 37). These pentapeptide products are thought to be potent activators of AmpR for the induction of *ampC* expression (13). Thus, *ampC* expression is a sensitive reporter for problems with PG crosslinking.

Beta-lactams covalently modify the TPase active sites of PBPs and inhibit PG crosslinking (22). These drugs do not block the GTase activity of the polymerase enzymes. Thus, uncrosslinked PG glycans are produced following drug treatment (23). In the related gram-negative bacterium *Escherichia coli*, these uncrosslinked strands have been shown to be rapidly degraded by a lytic transglycosylase (LT) enzyme (23). LTs cleave the glycan strand and generate disaccharide-peptide products with a 1,6-anhydro linkage on the MurNAc sugar (24). These so-called anhydro-muropeptides (AMPs) are produced by LTs during normal growth as these enzymes help promote the high turnover of the mature PG observed per generation (ca. 40%/generation) (**Fig. 1**) (25). In this case, the turnover products are primarily in the tetrapeptide form (26) because the terminal D-Ala of the stem peptide is either removed in the process of crosslinking or rapidly trimmed by carboxypeptidases. However, when nascent PG is processed by LTs during beta-lactam treatment, the AMPs produced are in the pentapeptide form (23). These are the AMPs that are most likely sensed by the AmpR regulator, converting it to an activator of *ampC* expression (**Fig. 1**) (13). Thus, *ampC* regulation is elegantly tuned to detect problems with nascent PG crosslinking as a proxy for the presence of beta-lactams.

To identify factors important for proper PG biogenesis, we have been using a fusion of the *lacZ* reporter gene to the *ampC* promoter (P*_ampC_*::*lacZ*) (27) to screen for mutants that aberrantly induce *ampC* expression in the absence of beta-lactams. This screen previously identified a missing enzyme in the PG recycling pathway of *P. aeruginosa* that is conserved in many gram-negative bacteria (28). Here, we report the surprising finding that defects in the LT enzyme SltB1 aberrantly induce *ampC*. LTs are typically associated with the production of the *ampC* inducing signal. It is therefore counterintuitive that a defect in an LT would promote elevated *ampC* expression. We show that this induction requires MltG, an LT enzyme previously implicated in the turnover of nascent PG and the induction of *ampC* following beta-lactam treatment in *P. aeruginosa* (29–31). Our results therefore support a model in which SltB1 facilitates the insertion and crosslinking of new PG into the matrix and that its inactivation causes elevated turnover of nascent PG strands. Thus, LTs may also function as space making enzymes similar to endopeptidases that cleave cell wall crosslinks (18–21).

## RESULTS

### Inactivation of sltB1 induces ampC expression and promotes beta-lactam resistance

A wild-type *P. aeruginosa* strain (PAO1) carrying a P*_ampC_::lacZ* reporter (27) inserted at the *attB* locus was mutagenized with a transposon. The resulting mutant library was plated on agar containing X-gal to identify insertions causing *ampC* expression in the absence of beta-lactam treatment. Prior screens using this library identified transposon mutants in the *mupP* gene, which led to its functional characterization as an enzyme important for the recycling of PG fragments (28). We continued to screen the library and identified an additional mutant forming solid-blue colonies on X-gal agar, a phenotype indicative of aberrant *ampC* induction. PCR-based mapping revealed that this isolate had a transposon inserted in the *sltB1* (*PA4001*) gene, encoding the LT enzyme SltB1.

A deletion mutation of *sltB1* was previously found to promote beta-lactam resistance (32, 33). However, in these studies, the authors did not detect elevated AmpC production in an *sltB1* mutant by nitrocefin hydrolysis assays or immunoblotting despite observing an *ampC* requirement for the resistance phenotype (32, 33). It was therefore concluded that beta-lactam resistance was not due to *ampC* induction but instead was likely to result from the inactivation of a lysis pathway involving cell wall damage caused by SltB1 (32, 33). The identification of the transposon insertion in *sltB1* in our screen argues against this interpretation and for a more direct role of SltB1 inactivation in *ampC* induction.

To validate the results from the screen, an in-frame deletion of *sltB1* similar to the previously published deletion (32, 33) was generated in the reporter strain. An aliquot of the mutant culture was spotted on agar containing X-gal alongside cultures of wild-type, a Δ*dacB* mutant known to promote high-level *ampC* induction, and a mutant lacking *ampR* that is defective for *ampC* expression (8, 34). As expected, the spots from the wild-type and Δ*ampR* culture remained white whereas that from the Δ*dacB* culture turned blue (**Fig. 2A**). The spot from the Δ*sltB1* culture also turned blue, indicating that SltB1 inactivation induces the *ampC* promoter in the absence of beta-lactams (**Fig. 2A**). Quantification of beta-galactosidase activity confirmed that the P*_ampC_::lacZ* reporter was induced in the Δ*sltB1* strain relative to wild-type (**Fig. 2B**). Thus, deletion of *sltB1* induces expression from the *ampC* promoter.

**Figure 2.**
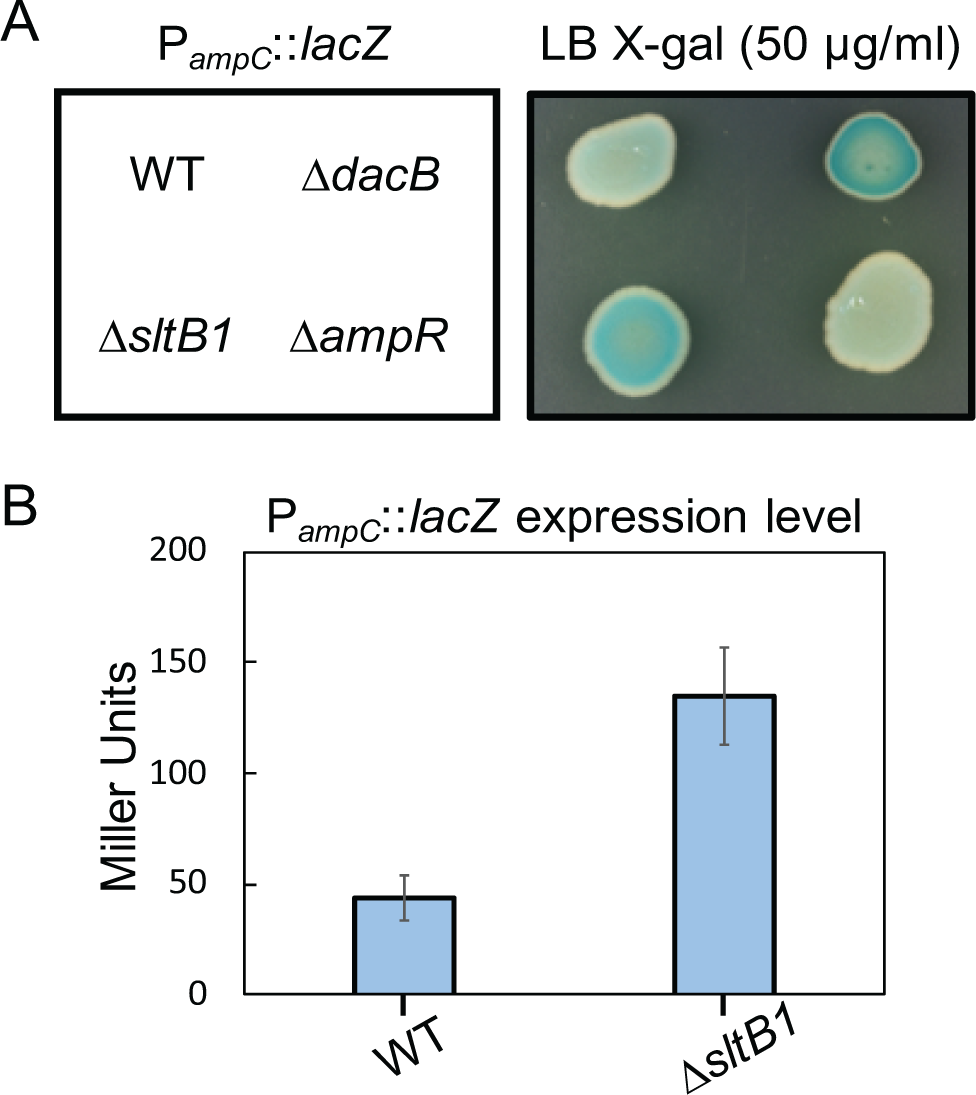
P*_ampC_*::*lacZ* is induced in an *sltB1* deletion strain. **(A)** Culture aliquots (5 µl) of strains CF262 [PAO1 WT], CF268 [Δ*dacB*], CF1143 [Δ*sltB1*], and CF604 [Δ*ampR*] containing the P*_ampC_*::*lacZ* reporter were spotted onto LB agar containing X-gal (50 µg/ml), grown overnight at 30°C, and photographed. **(B)** β-galactosidase activity in Miller Units was measured in liquid cultures of the indicated strains. Results shown are the average of 3 assays with 2 or 3 biological replicates per strain and the error bars represent the standard deviation.

To monitor the effect of SltB1 inactivation on native P*_ampC_* induction, we tested the beta-lactam resistance of mutant cells and measured the production of AmpC using a nitrocefin hydrolysis assay. Consistent with prior results, the Δ*sltB1* strain was resistant to the antipseudomonal beta-lactams ceftazidime and piperacillin, as was a control Δ*dacB* strain (**Fig. 3A**). Similarly, and in contrast to previous work (32), a Δ*sltB1* mutant with an empty vector showed elevated AmpC activity in the nitrocefin assay relative to wild-type cells (**Fig. 4A**). Deletion of *sltB1* did not strongly affect the induction of AmpC activity by the beta-lactam cefoxitin (**Fig. 4B**). The *sltB1* gene is in a putative operon that includes several genes encoding PG synthesis and remodeling proteins, including the downstream *rlpA* gene that also encodes an LT enzyme (**Fig. 3B**) (35). Notably, normal beta-lactam sensitivity and AmpC activity was restored to Δ*sltB1* cells upon expression of *sltB1* from a plasmid (**Fig. 3C and 4A**), indicating that the phenotype was caused by the inactivation of SltB1 and not an effect of the deletion on the expression of nearby genes.

**Figure 3.**
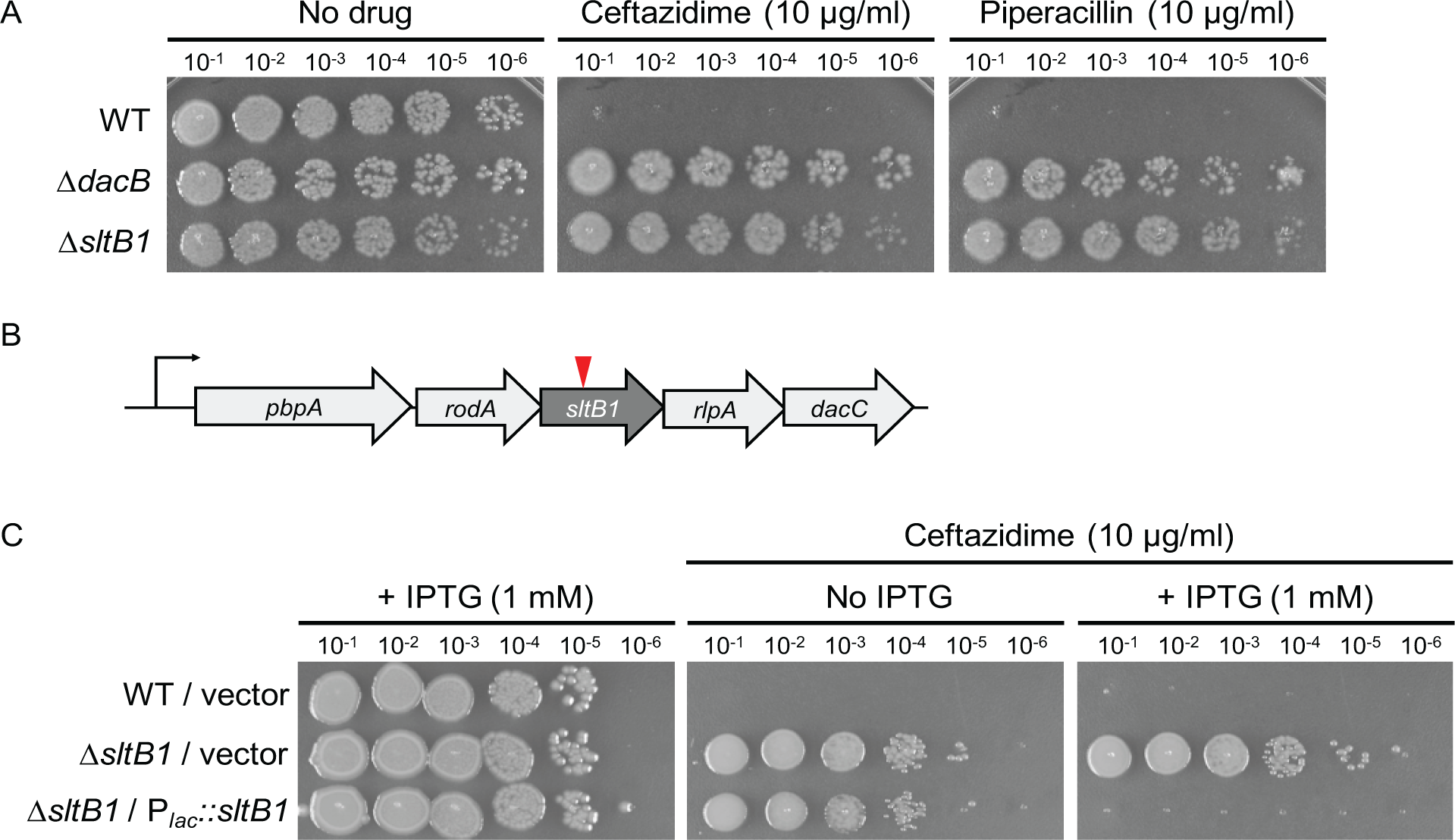
Inactivation of SltB1 promotes beta-lactam resistance. **(A)** Cultures of strains PAO1 [WT], CF155 [Δ*dacB*], and CF1105 [Δ*sltB1*] were serially diluted and 5 µl of each dilution was spotted onto LB agar supplemented with ceftazidime (10 µg/ml) or piperacillin (10 µg/ml) as indicated. **(B)** Diagram of the genetic locus harboring *sltB1*. The *pbpA* and *rodA* genes encode the RodA-PBP2 complex that forms the essential PG synthase of the cell elongation system (14). The *rlpA* gene encodes an LT enzyme that functions in cell division by promoting daughter cell separation (35), and *dacC* encodes a carboxypeptidase that trims PG peptides (49). (**C**) Cultures of the strains in (**A**) harboring an empty vector (pKHT103) or a plasmid (pCF533) encoding *sltB1* under control of the lac promoter (P*_lac_*) were diluted and plated as in panel (**A**) on media with or without IPTG inducer and/or ceftazidime (10 µg/ml) as indicated.

**Figure 4.**
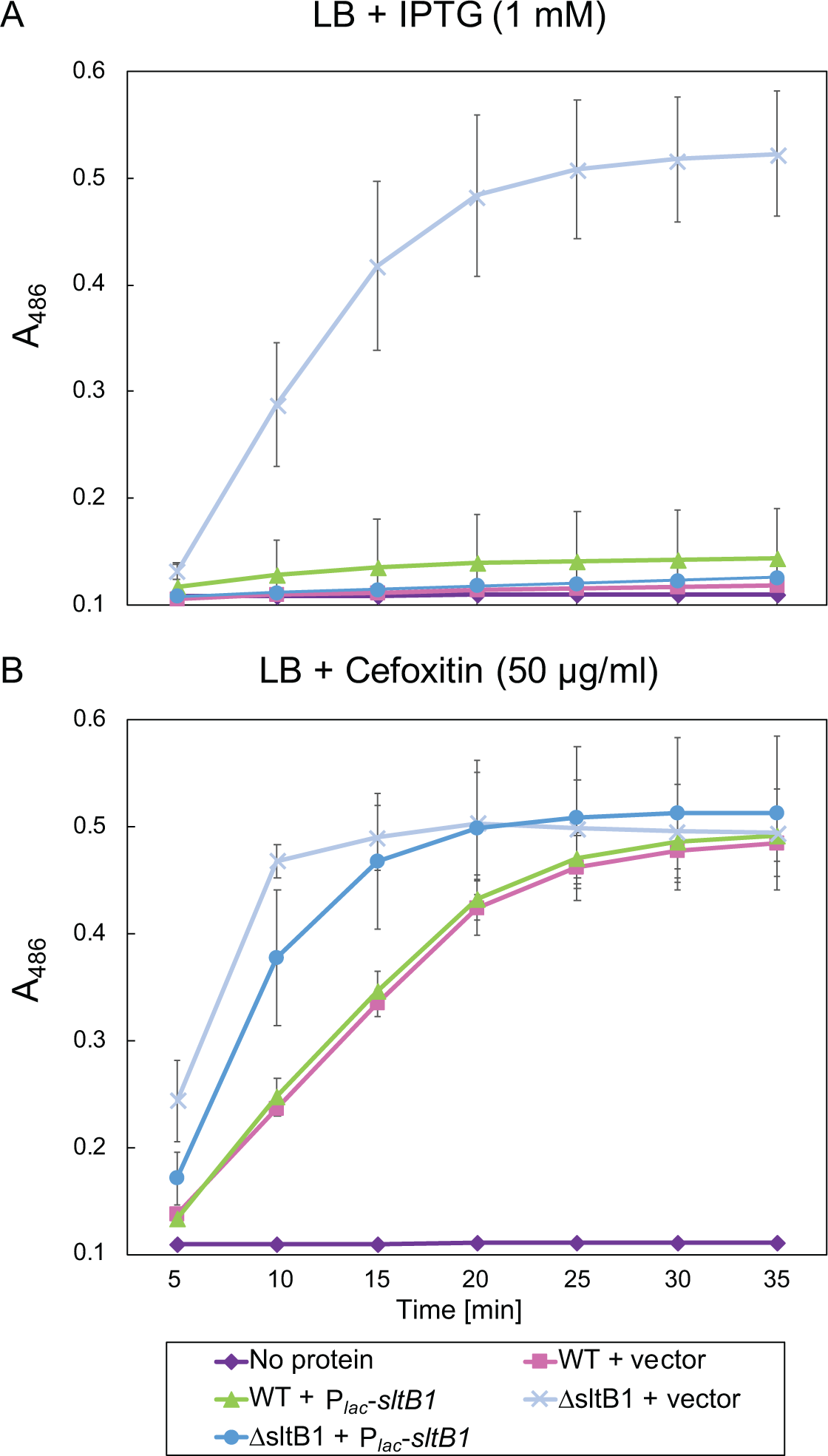
AmpC activity is elevated upon inactivation of SltB1. Assay of nitrocefin hydrolysis by cells of PAO1 [WT] or CF1105 [Δ*sltB1*] with plasmids pKHT103 [vector control] or pCF533 [P*_lac_*::*sltB1*] as indicated. Cells were grown in LB with IPTG (1 mM) or the *ampC* inducer cefoxitin (50 µg/ml) as indicated. Data are the mean of three independent assays each for two biological replicates with the error bars indicating the standard error.

### Requirements for ampC induction in ΔsltB1 cells

The results presented thus far suggest that mutants lacking SltB1 are resistant to beta-lactam treatment via the induction of *ampC* expression as opposed to alternative mechanisms proposed previously (32, 33). To further test this possibility, we investigated the requirements for resistance in the Δ*sltB1* background. Deletion of *ampC* or the *ampR* gene encoding the *ampC* transcriptional activator eliminated the piperacillin resistance phenotype of Δ*sltB1* cells and resulted in the loss of AmpC accumulation as determined by immunoblot (**Fig. 5A-B**). Beta-lactam resistance and AmpC accumulation were also blocked in the Δ*sltB1* mutant by inactivation of AmpG, the transporter that imports the AMP products from cell wall degradation that are sensed by AmpR to activate *ampC* expression (**Fig. 1 and 5A-B**) (36, 37). We therefore conclude that *ampC* is being induced in the Δ*sltB1* cells by the canonical induction mechanism.

**Figure 5.**
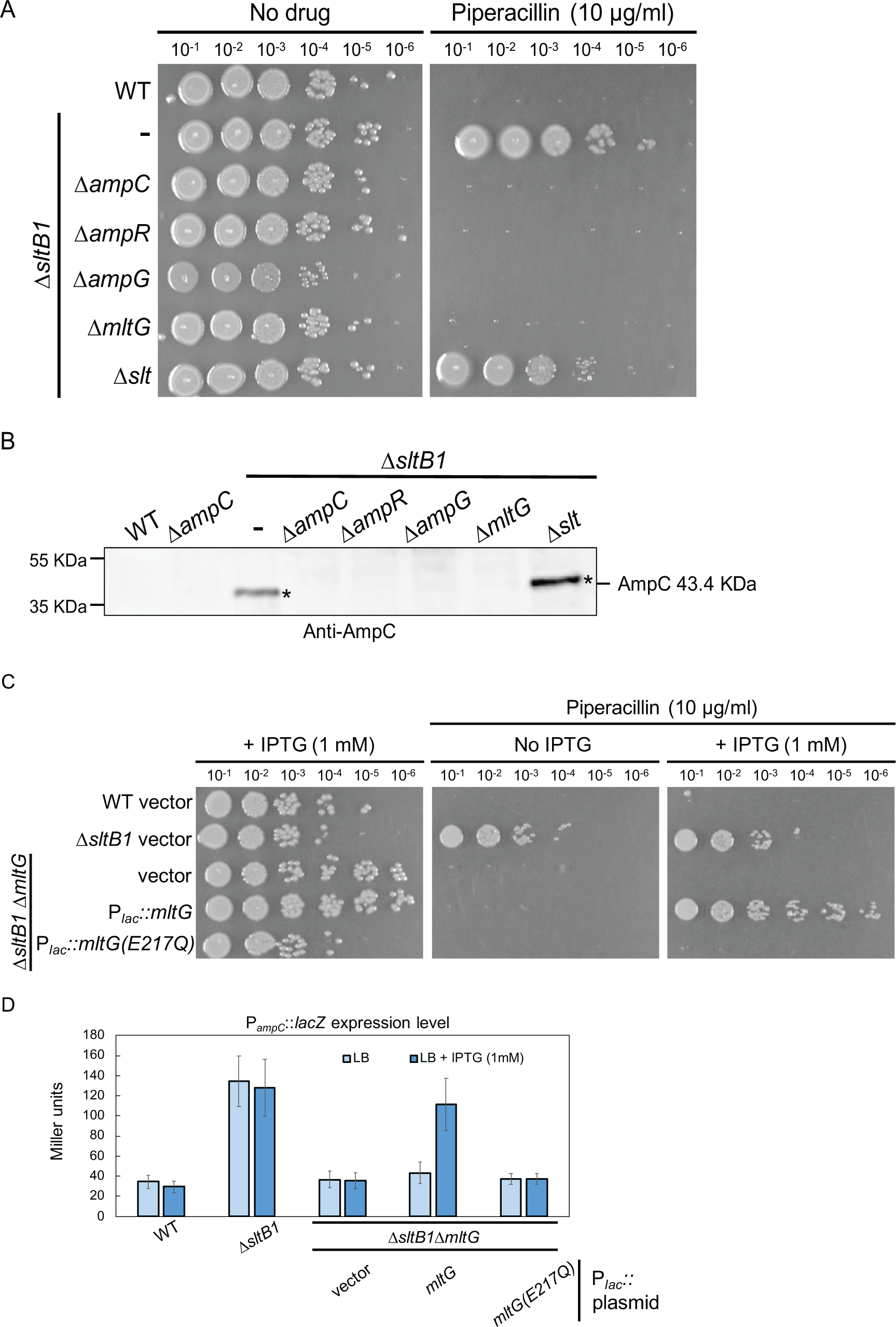
Requirements for beta-lactam resistance and AmpC induction in Δ*sltB1* cells. **(A)** Cultures of strains PAO1 [WT], CF1105 [Δ*sltB1*], CF368 [Δ*sltB1* Δ*ampC*], CF370 [Δ*sltB1*Δ*ampR*], CF372 [Δ*sltB1* Δ*ampG*], CF1416 [Δ*sltB1* Δ*mltG*] and CF378 [Δ*sltB1* Δ*slt*] were serially diluted and 5 µl of each dilution was spotted onto LB agar with or without piperacillin (10 µg/ml) as indicated. (**B**) Immunoblot for AmpC protein using the strains from panel (**A**). (**C**) Cultures of strains PAO1 [WT], CF1105 [Δ*sltB1*], and CF1416 [Δ*sltB1* Δ*mltG*] containing plasmids pKHT103 [vector control], pCF658 [P*_lac_*::*mltG*], or pCF1328 [P*_lac_*::*mltG(E217Q)*] were serially diluted and 5 µl of each dilution was spotted onto LB agar with or without piperacillin (10 µg/ml) and/or IPTG (1 mM) as indicated. (**D**) Derivatives of the strains in (**C**) with the P*_ampC_*::*lacZ* reporter were assayed for β-galactosidase activity as in Figure 2B.

In *E. coli*, uncrosslinked PG strands are generated upon beta-lactam treatment, and these strands are rapidly degraded by the LT enzymes Slt and MltG, with Slt playing the predominant role (23, 29). The action of these LTs generates the AMP products that in *P. aeruginosa* would serve as inducers of *ampC* expression. We therefore investigated whether Slt or MltG are required for *ampC* induction in the Δ*sltB1* background. Deletion of *slt* did not have a strong effect on the piperacillin resistance of Δ*sltB1* cells and actually appeared to increase AmpC production based on immunoblotting (**Fig. 5-A-B**). By contrast, inactivation of MltG restored piperacillin sensitivity to the Δ*sltB1* mutant and resulted in AmpC being undetectable in these cells (**Fig. 5A-B**). The piperacillin sensitivity of Δ*sltB1* Δ*mltG* cells was converted back to resistance upon expression of *mltG*(WT) from a plasmid but not by *mltG*(E217Q) encoding a catalytically dead MltG enzyme (**Fig. 5C**). Similarly, the induction of the P*_ampC_*::*lacZ* reporter in Δ*sltB1* Δ*mltG* cells was restored by plasmid-based production of MltG(WT) but not MltG(E217Q) (**Fig 5D**). These results are consistent with PG crosslinking being partially defective in the Δ*sltB1* mutant, resulting in the production of uncrosslinked strands that are turned over by MltG to produce the AMP products that induce *ampC* expression.

### Overexpression of catalytically inactivated SltB1 induces ampC expression

PG cleaving enzymes have been proposed to function as part of multi-protein complexes that help coordinate their activities with those of PG synthases (38). Notably, SltB1 has been found to interact with PBP2 (39, 40), a bPBP involved in cell elongation and shape determination (17). We therefore wondered whether SltB1 might have a limited number of binding sitesor binding partners in the cell that are required for its function. To test this possibility, we overproduced a FLAG-tagged variant of either SltB1(WT) or SltB1(E135A), a catalytically inactive variant, in wild-type cells and monitored beta-lactam resistance and AmpC production. Overexpression of *sltB1(WT)-FLAG* did not alter the ceftazidime sensitivity of the wild-type strain nor did it lead to the detectable induction of AmpC production (**Fig. 6**). However, overproduction of SltB1(E135A)-FLAG to levels equivalent to that of the wild-type protein conferred a ceftazidime resistance phenotype to otherwise wild-type cells and led to AmpC production (**Fig. 6**). This resistance phenotype required *ampC*, *ampR*, and *mltG* similar to that of the *sltB1* deletion (**Fig. 6**). We therefore conclude that SltB1 is likely to act from a finite set of binding sites in the cell, possibly in the form of interaction sites with partner PG synthases like PBP2.

**Figure 6.**
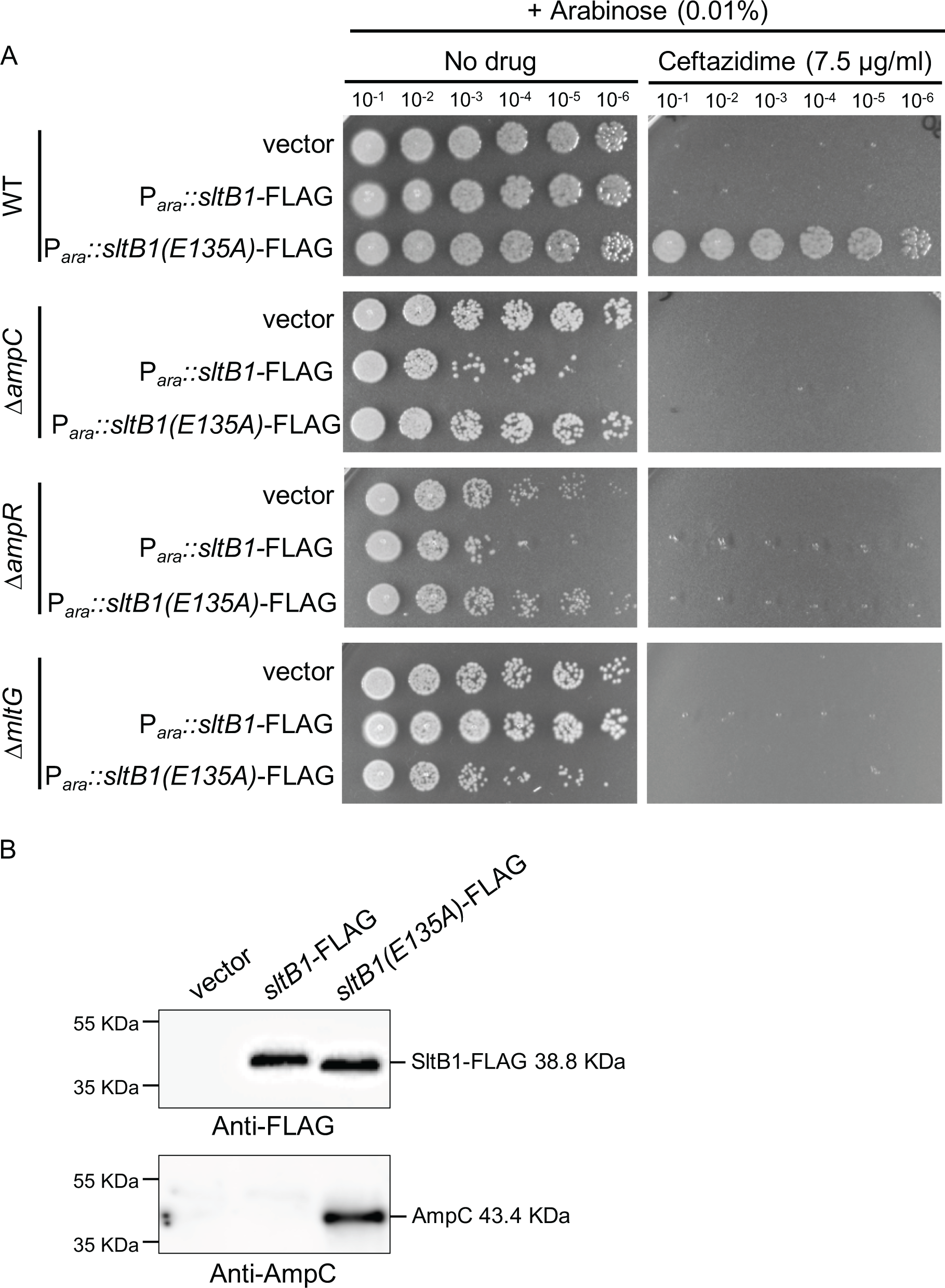
Overproduction of catalytically inactive SltB1 induces AmpC production and promotes beta-lactam resistance. (**A**). Cultures of strain PAO1 [WT], CF 612 [Δ*ampC*], CF550 [Δ*ampR*] and CF1410 [Δ*mltG*] with plasmids pJN105 [vector control], pCF1009 [P*_ara_*::*sltB1-FLAG*], or pCF1010 [P*_ara_*::*sltB1(E135A)-FLAG*] were serially diluted and 5 µl of each dilution was spotted onto LB agar supplemented with arabinose (0.01%) and ceftazidime (7.5 µg/ml) as indicated. (**B**) Immunoblots for FLAG-tagged proteins and AmpC protein using the wild-type strains from panel (**A**).

## DISCUSSION

Mutations that inactivate LT enzymes have been found to increase the expression of beta-lactamase genes in several different gram-negative organisms (32, 33, 41, 42). However, the mechanism(s) behind this phenomenon and how it relates to the role(s) of LTs in cell wall biogenesis has remained unclear. It was originally reported that mutants defective for SltB1 in *P. aeruginosa* confer elevated beta-lactam resistance via the inactivation of a cell death pathway involving this PG cleaving enzyme (32, 33). This mechanism was proposed because the authors did not detect elevated AmpC activity or protein levels in extracts from Δ*sltB1* cells despite the resistance phenotype being dependent on *ampC*. By contrast, our results indicate that Δ*sltB1* cells are induced for *ampC* expression, which was shown in several ways: (i) via a P*_ampC_*::*lacZ* reporter gene, (ii) AmpC activity assays, and (iii) immunoblotting for AmpC. Furthermore, induction was shown to depend on the importer AmpG that transports AMP products into the cytoplasm and the AmpR activator that stimulates *ampC* expression when it senses these molecules (34, 36, 37). Why the previous reports did not also observe *ampC* induction is not known. Nevertheless, the results presented here strongly support the conclusion that SltB1 inactivation confers beta-lactam resistance by activating *ampC* expression via the canonical mechanism involving the sensing of AMP turnover products.

An important clue to the mechanism by which SltB1 inactivation causes *ampC* induction came from the observation that it requires another LT enzyme called MltG. In *E. coli*, this inner membrane anchored LT has been implicated in the cleavage of nascent PG glycans, potentially functioning to sever their connection with the membrane and associated PG synthase thereby terminating their synthesis (29). In *P. aeruginosa*, MltG was found to be a key target of bulgecin A, an LT inhibitor that sensitizes *P. aeruginosa* and other gram-negative bacteria to beta-lactams (31). The implication of this finding is that MltG is likely involved in the turnover of uncrosslinked strands produced upon beta-lactam treatment, preventing the toxic side-effects of these glycans (23) and providing the AMP products for *ampC* induction. Accordingly, MltG was shown to be required for nascent PG turnover following cefsulodin treatment of *E. coli* (29). Notably, MltG was also recently shown to be required for *ampC* induction and beta-lactam resistance in a clinical isolate of *P. aeruginosa* (30). Therefore, based on the MltG requirement for *ampC* activation in Δ*sltB1* cells, we infer that the inactivation of SltB1 causes an increase in the formation of uncrosslinked nascent PG strands that are degraded by MltG to produce the AMP signals that induce beta-lactamase expression.

How might an LT enzyme like SltB1 that cuts glycans prevent the formation of uncrosslinked PG strands? One attractive explanation comes from the observed interaction between PBP2 and SltB1 (39, 40). PBP2 is a bPBP with TPase activity that together with the SEDS GTase RodA forms the essential cell wall synthase of the Rod system (elongasome) responsible for cell elongation and shape determination (14–17). Although the physiological relevance of the SltB1-PBP2 interaction has yet to be demonstrated, the location of the *sltB1* gene just downstream of the genes encoding RodA and PBP2 (**Fig. 3B**) suggests that SltB1 may be a non-essential component of the Rod system that supports its function. We therefore propose that SltB1 does so by associating with PBP2 and/or other PG sythases, positioning the LT enzyme to cut glycan strands in the mature PG matrix that may interfere with the insertion of new PG material during cell wall expansion. Thus, in the absence of SltB1, these problematic glycans remain, resulting in increased production of uncrosslinked nascent PG glycans that are degraded by MltG to generate the *ampC* inducing signal. Consistent with this model, catalytically inactive SltB1 exerts a dominant-negative Δ*sltB1-*like phenotype indicating that SltB1 has a limited number of binding sites in the cell that are required for its function.

In conclusion, we have used an *ampC* induction phenotype in *P. aeruginosa* to clarify the mechanism by which mutations in *sltB1* confer beta-lactam resistance. Our results also revealed a potential function for LT enzymes as space makers promoting the insertion of new PG strands into the mature wall. Such a function is commonly only ascribed to endopeptidases that cut PG crosslinks (14, 18–21). However, the PG matrix is unlikely to be so neatly arranged that crosslinks are the only impediment to the insertion of nascent glycans. Active synthases are also likely to encounter mature glycans that cross their path and require removal for the new strand to be effectively incorporated. Like SltB1, the inactivation of other LTs has also been associated with beta-lactamase induction (30, 41, 42), suggesting that a subset of the many LTs encoded by *P. aeruginosa* and other gram-negative bacteria may function in a space making capacity.

## METHODS AND MATERIALS

### Media, bacterial strains, and plasmids

*P. aeruginosa* PAO1 cells were grown in LB (1% tryptone, 0.5% yeast extract, 0.5% NaCl). As indicated, the medium was supplemented with 1 mM IPTG (isopropyl β-D-1-thiogalactopyranoside), 0.01% arabinose, 5% sucrose, or 50 µg/ml X-gal (5-bromo-4-chloro-3-indolyl-beta-D-galactopyranoside). For plasmid maintenance or integration, gentamicin (Gm), tetracycline (Tet), or carbenicillin (Carb) were used at a concentration of 30, 50 and 200 μg/mL, respectively. For AmpC beta-lactamase induction, cefoxitin (Fox) was used at a concentration of 50 μg/mL. Unless otherwise indicated, antipseudomonal antibiotics for viability/sensitivity assays were used at 10 μg/mL (piperacillin; Pip or ceftazidime; Caz). All *P. aeruginosa* strains used in the reported experiments are derivatives of PAO1. *E. coli* cells were grown in LB. For plasmid maintenance or selection, antibiotic concentration used was 10 μg/mL (gentamycin; Gm or Tetracycline; Tet). The bacterial strains and plasmids used in this study are listed in **Table S1, S2 and S3**. Detailed descriptions of the strain and plasmid construction procedures can be found in the Supplementary Material.

### *P. aeruginosa* viability assay

For viability assays, overnight cell cultures were normalized to an OD of 1 and subjected to serial 10-fold dilutions with LB. Five microliters of each dilution was then spotted onto the indicated agar and plates were incubated at 30°C for 24 hours prior to imaging.

### *P. aeruginosa* electroporation

*P. aeruginosa* strains were made competent using previously described methods (43). Briefly, 4 mL of overnight cultures grown at 37°C were centrifuged and washed twice with 1 mL 300 mM sucrose. Cell pellets were resuspended in 500 µL of 300 mM sucrose and 100 µL were used for electroporation. 1µL of replicative plasmid was used for the electroporation, using the following settings: 25 mF, 200 O, 2.5 kV. LB medium (1mL) was added and the cells incubated with shaking (200 rpm) for 1 h at 37 °C. Cells were then plated on the appropriate selective medium.

### Screen for mutants that induce *ampC* expression

The screening procedure was described previously (28). Briefly, *P. aeruginosa* strain CF263 [PAO1 (P*_ampC_::lacZ*)] was transposon mutagenized by mating with the *E. coli* donor SM10(λpir) harboring a mariner transposon delivery vector pIT2 (44). The transposon confers tetracycline resistance. Mating mixtures were plated on LB agar supplemented with tetracyline (50 µg/ml) to select for transposon mutants and nalidixic acid (25 µg/ml) to select against the *E. coli* donor. The resulting collection of colonies was resuspended in LB broth and stored at −80°C. Dilutions of the library were plated on LB containing X-gal (50 µg/ml) to identify mutants with a constitutively active P*_ampC_::lacZ* reporter.

### Mapping of transposon insertion sites

Transposon insertions were mapped using arbitrarily-primed PCR (44). Transposon-chromosomal junctions were amplified from mutant chromosomal DNA using the primers Rnd1-PA (5’-GGCCACGCGTCGACTAGTACNNNNNNNNNNGATAT 3’) and LacZ211 (5’-TGC GGG CCT CTT CGC TAT TA-3’). The resulting PCR reaction was used for a second PCR with primers Rnd2-PA (5’-GGCCACGCGTCGACTAGTAC-3’) and LacZ148 (5’-GGG TAA CGC CAG GGT TTT CC-3’). The final PCR product was sequenced using the transposon specific primer LacZ-124L (5’ CAG TCA CGA CGT TGT AAA ACG ACC). The transposon-chromosomal DNA junction was identified in the sequencing reads using a nucleotide BLAST search (45) against the PAO1 genome (46).

### β-galactosidase assay

β-Galactosidase assays were performed at room temperature. Cells from 100 μl of culture at OD_600_ = 0.1-0.6 were lysed with 30 μl of chloroform and mixed with 700 μl of Z buffer (60 mM Na_2_HPO_4_, 40 mM NaH_2_PO_4_, 10 mM KCl and 1 mM MgSO_4_ heptahydrate). Each reaction then received 200 μl of o-nitrophenyl-β-D-galactopyranoside (ONPG, 4 mg/ml in 0.1 M KPO_4_ pH7.0) and the reaction was timed. After a medium-yellow color developed, the reaction was stopped with 400 μl of 1M Na_2_CO_3_. The OD_420_ of the supernatant was determined and the units of activity (Miller Units) were calculated using the equation: U= (OD_420_ * 1000) / (OD_660_ * time (in min) * volume of culture (in ml)).

### AmpC beta-lactamase activity assay

AmpC activity was assessed using nitrocefin hydrolysis. Overnight bacterial cultures were subcultured 1:20 in 3 mL LB or 6 mL LB supplemented with 1mM IPTG and grown for 2 hr at 30°C and 200 rpm. The cultures were split 1:1 in 2 mL LB with or without 50 μg/mL cefoxitin (final concentration) and all cultures were incubated for an additional 1.5 hr at 30°C and 200 rpm. Following incubation, 1 mL of culture was pelleted at 2,300 x g for 5 minutes, washed once with 1 mL of 50 mM sodium phosphate buffer (pH 7.0) and resuspended in 1mL of the same cold buffer. Samples were placed on ice and lysed at 4°C by sonication with a microprobe (Q800R2, QSonica, Newtown, Connecticut, USA). Sonicated samples were centrifuged at 12,000 x g for 5 minutes at 4°C and supernatants were collected. The protein concentration was determined using a Bradford assay (47) with bovine serum albumin (BSA) as the standard (G-Biosciences, Geno technology inc., Saint-Louis, Missouri, USA). Nitrocefin hydrolysis assays were performed in 96-well plates. Each reaction had a final volume of 250 μl of 50 mM sodium phosphate buffer (pH 7.0) containing 10 μg of protein and 20 μg of nitrocefin (Thermo Fischer Scientific™ Oxoid, Waltham, Massachusetts, USA). Nitrocefin hydrolysis was monitored by measuring the absorbance at 486 nm every 5 minutes for 30 minutes at 30°C.

### Immunoblotting

Overnight bacterial cultures were subcultured 1:20 in 5 mL LB and grown for 4 hours. Bacteria were collected by centrifugation, washed once with Tris-HCl buffer (pH=8, 10 mM) and resuspended in 500 µL of the same cold buffer. The samples were then lysed at 4°C for 12 minutes at 60% amplitude with a pulse rate of 10 seconds ON/10 seconds OFF using a Qsonica sonicator. The samples were then centrifuged at 4°C at maximal speed to remove the cell debris. Supernatants were collected and a Bradford assay performed to measure the protein concentration. A total of 100 µg protein in a volume of 100µL were mixed with 100µL of 2X Laemmli buffer. Immunoblotting was performed by first separating 15 µL of each sample on 12% SDS-PAGE (<u>p</u>oly<u>a</u>crylamide <u>g</u>el <u>e</u>lectrophoresis) gels at 90V for 15 minutes and 120V for an hour. Proteins were transferred at 90V for an hour at 4°C to a 0.2 μm PolyVinylidene DiFluoride (PVDF) membranes (Whatman) previously soaked in methanol and rinsed with transfer buffer. Membranes were blocked using 5% (w/v) skim milk in Tris-Buffered Saline (10 mM Tris-HCl pH 7.5, 150 mM NaCl) supplemented with 0.1% (v/v) Tween-20 (TBS-T) for 1 hour. Membranes were incubated for 1 hour with α-AmpC primary antibody (1:1000 dilution in 5% skim milk in TBS-T, MyBioSource, MBS1493275, San Diego, USA) or α-FLAG primary antibody (1:1000 dilution in 5% skim milk in TBS-T, F7425, Sigma-Aldrich) at 4°C. The membranes were washed four times in TBS-T for 5 min each before incubation for 1 h with secondary antibody (anti-rabbit IgG HRP, 1:5000 dilution, Rockland 18-8816-33) in TBS-T with 5% (w/v) skim milk powder. The membranes were then washed four times with TBS-T for 5 min each before developing using SuperSignal West Pico PLUS Chemiluminescent Substrate (Thermo Fisher Scientific cat#34577) and imaged using the c600 Azure Biosystems platform.

## ACKNOWLEDGEMENTS

The authors would like to thank all members of the Bernhardt and Rudner labs for advice and helpful discussions. Special thanks to Stephen Lory and Simon Dove for help with *P. aeruginosa* methods and for providing strains and expert advice. This work was supported by the National Institute of Allergy and Infectious Diseases of the National Institutes of Health (R01 AI083365 and U19 AI158028 to T.G.B.) and Investigator funds from the Howard Hughes Medical Institute (T.G.B.). C.F. was supported in part by a postdoctoral fellowship from the Swiss National Science Foundation (project #P2GEP3_162073).

---

**Table S1.**
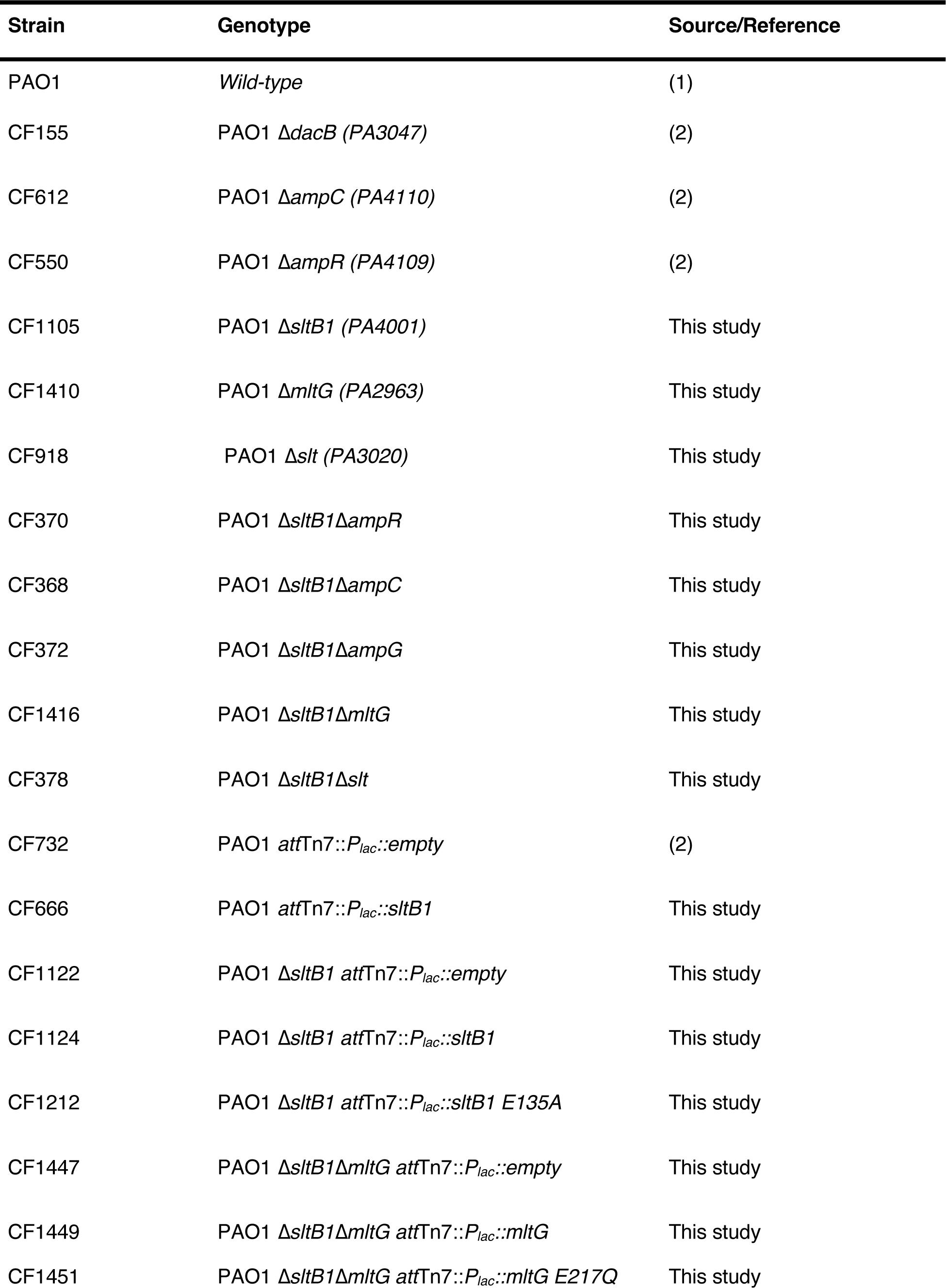
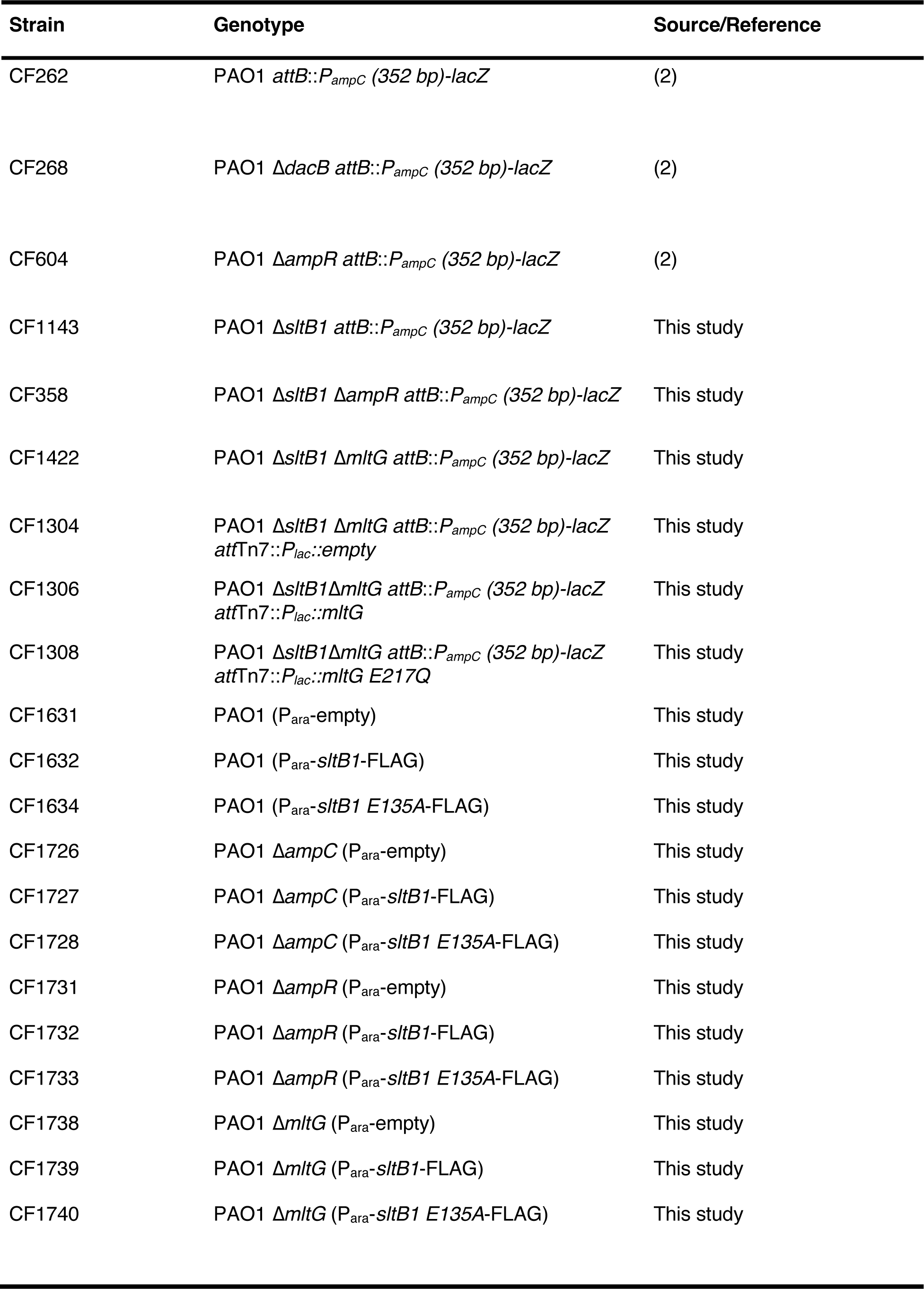
*Pseudomonas aeruginosa* strains used in this study.

| Strain | Genotype | Source/Reference |
| --- | --- | --- |
| PAO1 | *Wild-type* | (1) |
| CF155 | PAO1 $\Delta dacB$ (PA3047) | (2) |
| CF612 | PAO1 $\Delta ampC$ (PA4110) | (2) |
| CF550 | PAO1 $\Delta ampR$ (PA4109) | (2) |
| CF1105 | PAO1 $\Delta sltB1$ (PA4001) | This study |
| CF1410 | PAO1 $\Delta mltG$ (PA2963) | This study |
| CF918 | PAO1 $\Delta slt$ (PA3020) | This study |
| CF370 | PAO1 $\Delta sltB1\Delta ampR$ | This study |
| CF368 | PAO1 $\Delta sltB1\Delta ampC$ | This study |
| CF372 | PAO1 $\Delta sltB1\Delta ampG$ | This study |
| CF1416 | PAO1 $\Delta sltB1\Delta mltG$ | This study |
| CF378 | PAO1 $\Delta sltB1\Delta slt$ | This study |
| CF732 | PAO1 *attTn7::P<sub>lac</sub>::empty* | (2) |
| CF666 | PAO1 *attTn7::P<sub>lac</sub>::sltB1* | This study |
| CF1122 | PAO1 $\Delta sltB1$ *attTn7::P<sub>lac</sub>::empty* | This study |
| CF1124 | PAO1 $\Delta sltB1$ *attTn7::P<sub>lac</sub>::sltB1* | This study |
| CF1212 | PAO1 $\Delta sltB1$ *attTn7::P<sub>lac</sub>::sltB1 E135A* | This study |
| CF1447 | PAO1 $\Delta sltB1\Delta mltG$ *attTn7::P<sub>lac</sub>::empty* | This study |
| CF1449 | PAO1 $\Delta sltB1\Delta mltG$ *attTn7::P<sub>lac</sub>::mltG* | This study |
| CF1451 | PAO1 $\Delta sltB1\Delta mltG$ *attTn7::P<sub>lac</sub>::mltG E217Q* | This study |
| CF262 | PAO1 *attB::P<sub>ampC</sub> (352 bp)-lacZ* | (2) |
| CF268 | PAO1 Δ*dacB attB::P<sub>ampC</sub> (352 bp)-lacZ* | (2) |
| CF604 | PAO1 Δ*ampR attB::P<sub>ampC</sub> (352 bp)-lacZ* | (2) |
| CF1143 | PAO1 Δ*sltB1 attB::P<sub>ampC</sub> (352 bp)-lacZ* | This study |
| CF358 | PAO1 Δ*sltB1 ΔampR attB::P<sub>ampC</sub> (352 bp)-lacZ* | This study |
| CF1422 | PAO1 Δ*sltB1 ΔmltG attB::P<sub>ampC</sub> (352 bp)-lacZ* | This study |
| CF1304 | PAO1 Δ*sltB1 ΔmltG attB::P<sub>ampC</sub> (352 bp)-lacZ*<br>*attTn7::P<sub>lac</sub>::empty* | This study |
| CF1306 | PAO1 Δ*sltB1 ΔmltG attB::P<sub>ampC</sub> (352 bp)-lacZ*<br>*attTn7::P<sub>lac</sub>::mltG* | This study |
| CF1308 | PAO1 Δ*sltB1 ΔmltG attB::P<sub>ampC</sub> (352 bp)-lacZ*<br>*attTn7::P<sub>lac</sub>::mltG E217Q* | This study |
| CF1631 | PAO1 (*P<sub>ara</sub>-empty*) | This study |
| CF1632 | PAO1 (*P<sub>ara</sub>-sltB1-FLAG*) | This study |
| CF1634 | PAO1 (*P<sub>ara</sub>-sltB1 E135A-FLAG*) | This study |
| CF1726 | PAO1 Δ*ampC* (*P<sub>ara</sub>-empty*) | This study |
| CF1727 | PAO1 Δ*ampC* (*P<sub>ara</sub>-sltB1-FLAG*) | This study |
| CF1728 | PAO1 Δ*ampC* (*P<sub>ara</sub>-sltB1 E135A-FLAG*) | This study |
| CF1731 | PAO1 Δ*ampR* (*P<sub>ara</sub>-empty*) | This study |
| CF1732 | PAO1 Δ*ampR* (*P<sub>ara</sub>-sltB1-FLAG*) | This study |
| CF1733 | PAO1 Δ*ampR* (*P<sub>ara</sub>-sltB1 E135A-FLAG*) | This study |
| CF1738 | PAO1 Δ*mltG* (*P<sub>ara</sub>-empty*) | This study |
| CF1739 | PAO1 Δ*mltG* (*P<sub>ara</sub>-sltB1-FLAG*) | This study |
| CF1740 | PAO1 Δ*mltG* (*P<sub>ara</sub>-sltB1 E135A-FLAG*) | This study |

**Table S2.**
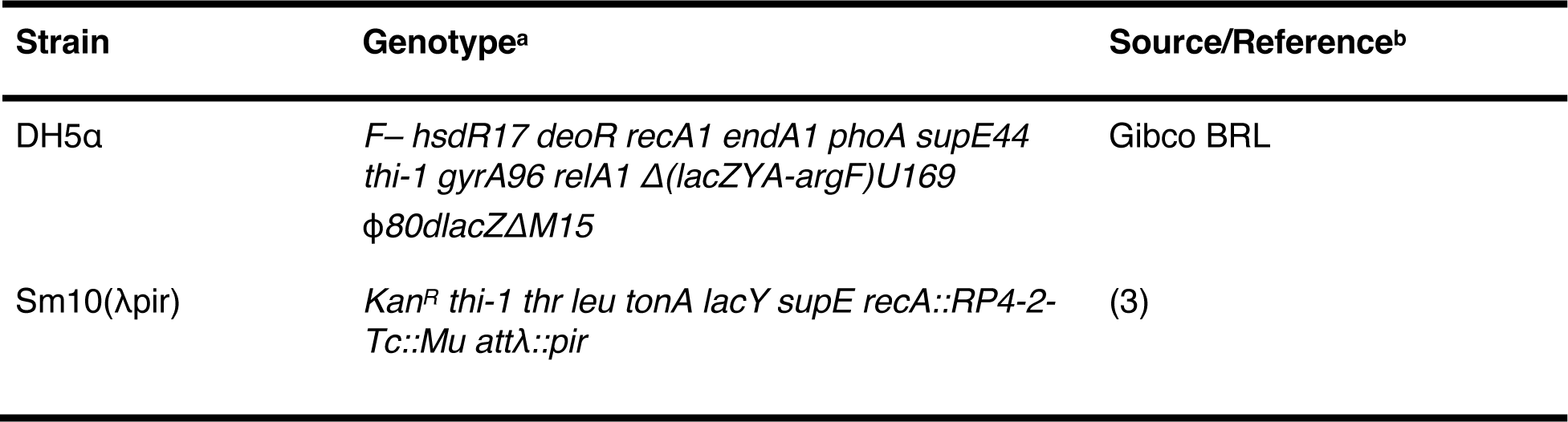
*Escherichia coli* strains used in this study.

**Table S3.**
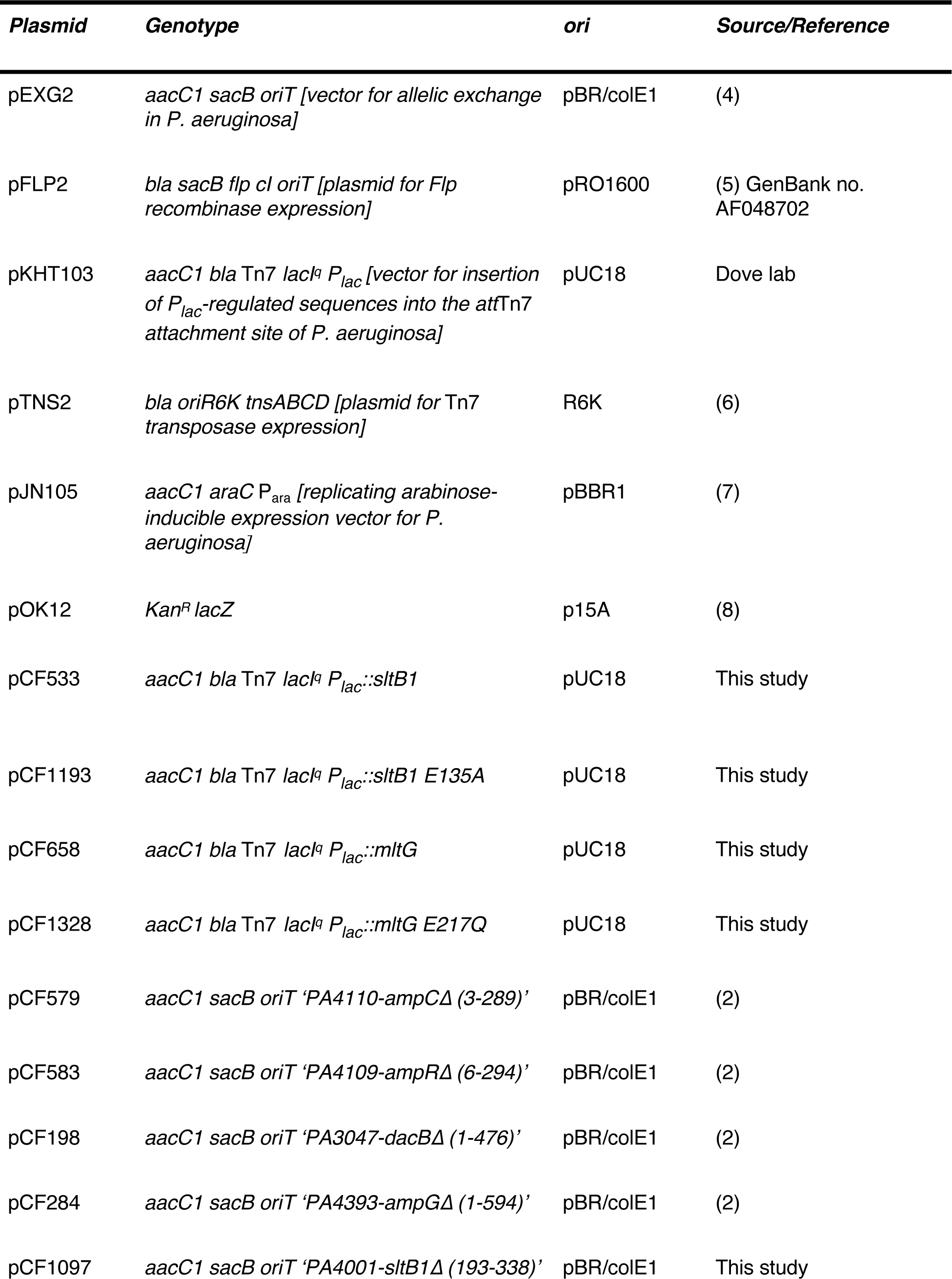
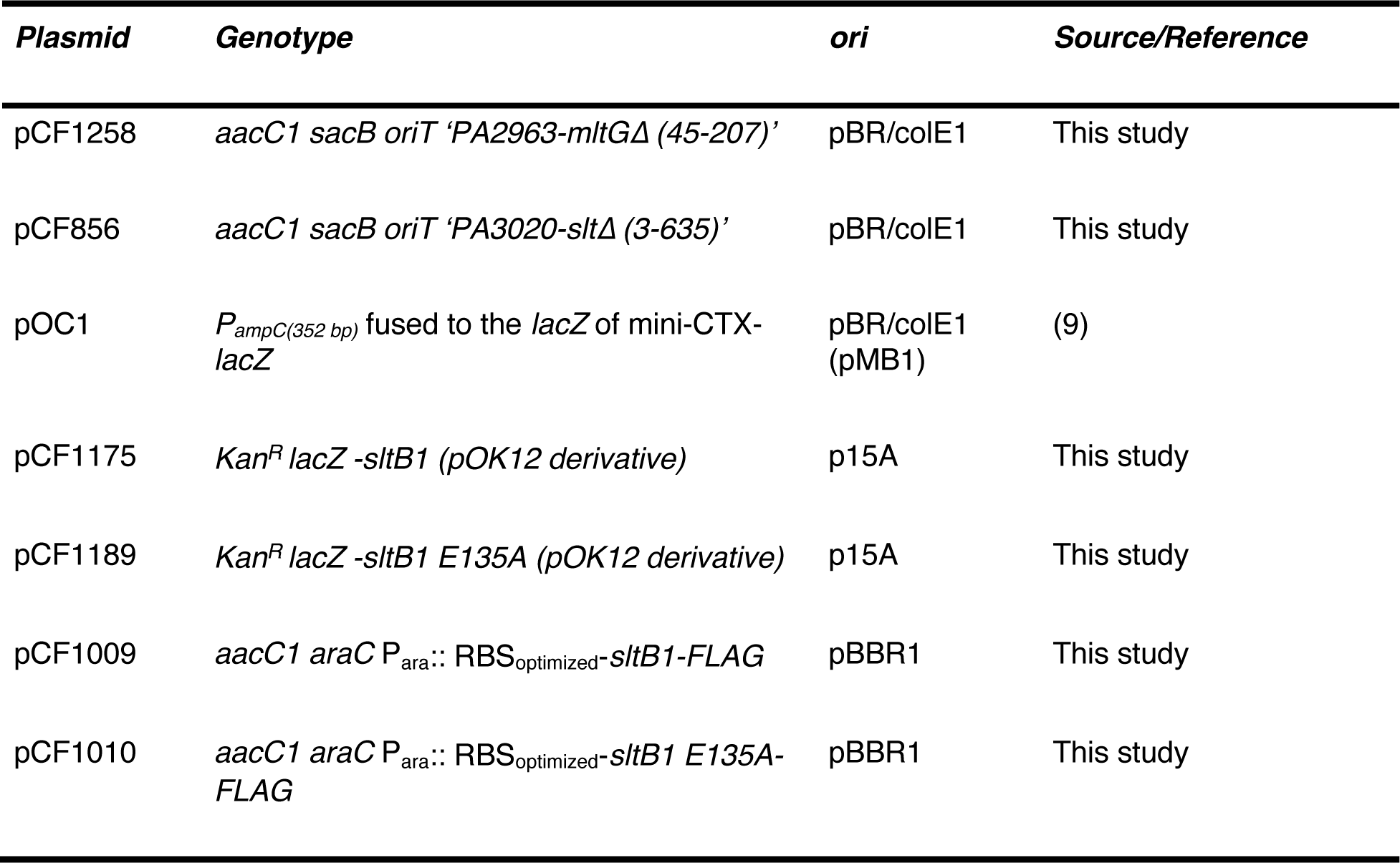
Plasmids used in this study.

| **Plasmid** | **Genotype** | **ori** | **Source/Reference** |
| --- | --- | --- | --- |
| pEXG2 | *aacC1 sacB oriT* [vector for allelic exchange in *P. aeruginosa*] | pBR/colE1 | (4) |
| pFLP2 | *bla sacB flp cl oriT* [plasmid for Flp recombinase expression] | pRO1600 | (5) GenBank no. AF048702 |
| pKHT103 | *aacC1 bla Tn7 lacI<sup>q</sup> P<sub>lac</sub>* [vector for insertion of *P<sub>lac</sub>*-regulated sequences into the attTn7 attachment site of *P. aeruginosa*] | pUC18 | Dove lab |
| pTNS2 | *bla oriR6K tnsABCD* [plasmid for Tn7 transposase expression] | R6K | (6) |
| pJN105 | *aacC1 araC P<sub>ara</sub>* [replicating arabinose-inducible expression vector for *P. aeruginosa*] | pBBR1 | (7) |
| pOK12 | *Kan<sup>R</sup> lacZ* | p15A | (8) |
| pCF533 | *aacC1 bla Tn7 lacI<sup>q</sup> P<sub>lac</sub>::sltB1* | pUC18 | This study |
| pCF1193 | *aacC1 bla Tn7 lacI<sup>q</sup> P<sub>lac</sub>::sltB1 E135A* | pUC18 | This study |
| pCF658 | *aacC1 bla Tn7 lacI<sup>q</sup> P<sub>lac</sub>::mltG* | pUC18 | This study |
| pCF1328 | *aacC1 bla Tn7 lacI<sup>q</sup> P<sub>lac</sub>::mltG E217Q* | pUC18 | This study |
| pCF579 | *aacC1 sacB oriT 'PA4110-ampCΔ (3-289)'* | pBR/colE1 | (2) |
| pCF583 | *aacC1 sacB oriT 'PA4109-ampRΔ (6-294)'* | pBR/colE1 | (2) |
| pCF198 | *aacC1 sacB oriT 'PA3047-dacBΔ (1-476)'* | pBR/colE1 | (2) |
| pCF284 | *aacC1 sacB oriT 'PA4393-ampGΔ (1-594)'* | pBR/colE1 | (2) |
| pCF1097 | *aacC1 sacB oriT 'PA4001-sltB1Δ (193-338)'* | pBR/colE1 | This study |
| pCF1258 | *aacC1 sacB oriT 'PA2963-mltGΔ (45-207)'* | pBR/colE1 | This study |
| pCF856 | *aacC1 sacB oriT 'PA3020-sltΔ (3-635)'* | pBR/colE1 | This study |
| pOC1 | *P<sub>ampC(352 bp)</sub> fused to the lacZ of mini-CTX-lacZ* | pBR/colE1 (pMB1) | (9) |
| pCF1175 | *Kan<sup>R</sup> lacZ -sltB1 (pOK12 derivative)* | p15A | This study |
| pCF1189 | *Kan<sup>R</sup> lacZ -sltB1 E135A (pOK12 derivative)* | p15A | This study |
| pCF1009 | *aacC1 araC P<sub>ara::</sub> RBS<sub>optimized</sub>-sltB1-FLAG* | pBBR1 | This study |
| pCF1010 | *aacC1 araC P<sub>ara::</sub> RBS<sub>optimized</sub>-sltB1 E135A-FLAG* | pBBR1 | This study |

## Plasmid construction

For all plasmid constructions (**Table S3**), PCR was performed using Phusion or the Q5 polymerase (New England Biolabs) according to the manufacturer’s instructions. Plasmid DNA was purified using the Zippy miniprep kits (Zymo Research) while PCR fragments were purified using a Qiaquick PCR purification kit (Qiagen). Unless otherwise indicated, plasmids were constructed using the Gibson isothermal assembly method (10) and the reactions were incubated at 50°C for 30 minutes.

To construct **pCF856** [*aacC1 sacB oriT ‘PA3020-sltΔ(3-635)*], which is used for deletion of *slt* (*PA3020*), the ∼800bp region upstream of *slt* was amplified from PAO1 genomic DNA (gDNA) using 5’-AAA TGT AAA GCA AGC TTC TGC AGG TCG ACT CTT TGG TCG GTT CGT CGA GCA GCA GCA GGT-3’ and 5’-TTG GGG GTC AGA AAT CGT CGA AGT AGC GTT CGC GCA TGA CTT ATC CGG GCA GGT GAT GT-3’ primers. The ∼800bp region downstream of the gene was amplified with 5’-ACA TCA CCT GCC CGG ATA AGT CAT GCG CGA ACG CTA CTT CGA CGA TTT CTG ACC CCC AA-3’ and 5’-GAA TTC GAG CTC GAG CCC GGG GAT CCT CTA TCG AGT TCG TCG TGG CCA AGC CCT GTT-3’ primers. The two resulting fragments were then combined by isothermal assembly into pEXG2, which was previously digested with XbaI.

To construct **pCF1097** [*aacC1 sacB oriT ‘PA4001-sltB1Δ(193-338)’*], which is used for deletion of *sltB1* (*PA4001*), the ∼800bp region upstream of *sltB1* was amplified from PAO1 genomic DNA (gDNA) using 5’-AAA TGT AAA GCA AGC TTC TGC AGG TCG ACT CTA GTT CAT GAA GCT GCT GAT GCC GAT GA-3’ and 5’-ATC AGG GAG GAG CGG ACA CGC TTG CTC AAT GGG CCG CCG GCG TAG GAG CCG GTC AGG CTG A-3’ primers. The ∼800bp region downstream of the gene was amplified with 5’-TCA GCC TGA CCG GCT CCT ACG CCG GCG GCC CAT TGA GCA AGC GTG TCC GCT CCT CCC TGA T-3’ and 5’-GAA TTC GAG CTC GAG CCC GGG GAT CCT TTT TTT TTT GAG TCG ATC TGC ACC GGC AGC-3’ primers. The two resulting fragments were then combined by isothermal assembly into pEXG2, which was previously digested with XbaI.

To construct **pCF1258** [*aacC1 sacB oriT ‘PA2963-mltGΔ(45-207)’*], which is used for deletion of *mltG* (*PA2963*), the ∼800bp region upstream of *mltG* was amplified from PAO1 genomic DNA (gDNA) using 5’-AAA TGT AAA GCA AGC TTC TGC AGG TCG ACT CTA CTG GCC TAT GGC GAC GGG CTG TTC GAG A-3’ and 5’-CTT TTC CAC CAG CGA GGC CAT GAT CAG AAC GTC CAG CAG GCG CTC CTC GGT CAA TTG-3’ primers. The ∼800bp region downstream of the gene was amplified with 5’-CAA TTG ACC GAG GAG CGC CTG CTG GAC GTT CTG ATC ATG GCC TCG CTG GTG GAA AAG-3’ and 5’-GAA TTC GAG CTC GAG CCC GGG GAT CCT AAT GCT CGG CCA GGG CAC GCT TAC CGA T-3’ primers. The two resulting fragments were then combined by isothermal assembly into pEXG2, which was previously digested with XbaI.

**pCF533** [*aacC1 bla Tn7 lacI^q^* P*_lac_*::*sltB1*]: For the construction of pCF533 for integration of an IPTG-inducible version of *sltB1* at the Tn7 attachment (*att*Tn7) site, *sltB1* was amplified from PAO1 genomic DNA using 5’-AGC TTA GTC GAC AGC TAG CCG GAT CCC CGG <u>GAG GAG</u> <u>GAT ACA T</u>GT GAA GAA CGC AAT GCA AGT ACT GCG TAC-3’ and 5’-AAG GGG TTA TGC TAA AGC TTG CAT GCG GTA CTC AAT GGG CAC CTC GCG CGC GGG CAA TCT-3’ primers. The optimized RBS is underlined. The resulting fragment was then inserted by Gibson isothermal assembly into a KpnI-digested expression vector pKHT103 to generate pCF533.

**pCF658** [*aacC1 bla Tn7 lacI^q^* P*_lac_*::*mltG*]: For the construction of pCF658 for integration of an IPTG-inducible version of *mltG* at the Tn7 attachment (*att*Tn7) site, *mltG* was amplified from PAO1 genomic DNA using 5’-AGC TTA GTC GAC AGC TAG CCG GAT CCC CGG <u>GAG GAG</u> <u>GAT ACA T</u>AT GCG CAA ACT GCT GGT GCT GCT GGA GAG-3’ and 5’-AAG GGG TTA TGC TAA AGC TTG CAT GCG GTA CTC ATT GTG GCG GCG GGG TGA TGG GC-3’ primers. The optimized RBS is underlined. The resulting fragment was then inserted by Gibson isothermal assembly into a KpnI-digested expression vector pKHT103 to generate pCF658.

Site-directed mutagenesis was performed using the QuikChange method (Stratagene) or PCR site directed mutagenesis.

For **pCF1175** [*Kan^R^ lacZ -sltB1* (pOK12 derivative)], the *sltB1* gene was amplified from wild-type genomic DNA using the primers *sltB1* HindIII 5’ optimized RBS (5’-AAA <u>AAG CTT</u> **GAG GAG GAT ACA T**GT GAA GAA CGC AAT GCA AGT ACT GCG TAC −3’) and *sltB1* XbaI 3’ (5’-AAA <u>TCT AGA</u> TCA ATG GGC ACC TCG CGC GCG GGC AAT CT −3’). The restriction sites are underlined and the optimized RBS is bolded. The PCR product was then digested with XbaI and HindIII and ligated into XbaI/HindIII digested pOK12. The primer sequences used for the mutagenesis are *sltB1 E135A* #1 (5’-ATC ATC GGC GTG <u>GCA</u> ACC TTC TTC GGC −3’) and *sltB1 E135A* #2 (5’-GCC GAA GAA GGT <u>TGC</u> CAC GCC GAT GAT −3’). The PCR was performed using KOD polymerase (Novagen) according to the manufacturer’s instructions (65°C annealing temperature and 2.5 minutes of extension for 20 cycles). The PCR product was directly treated for 5 hours at 37°C with 1 µl of DpnI restriction enzyme to digest the parental double-stranded DNA. A portion of the reaction (5 µl) was used to transform chemo-competent DH5α and transformants were selected on LB plates containing 25 µg/ml kanamycin. The plasmid with the correct mutation was identified by sequencing and was designated **pCF1189** [*Kan^R^ lacZ-sltB1 E135A* (pOK12 derivative)].

**pCF1193** [*aacC1 bla Tn7 lacI^q^ P_lac_::sltB1 E135A*]. For the construction of pCF1193 for integration of an IPTG-inducible version of *sltB1(E135A)* at the Tn7 attachment (*att*Tn7) site, *sltB1(E135A)* was amplified from pCF1189 [*Kan^R^ lacZ-sltB1 E135A* (pOK12 derivative)] plasmid DNA using 5’-AGC TTA GTC GAC AGC TAG CCG GAT CCC CGG <u>GAG GAG GAT ACA T</u>GT GAA GAA CGC AAT GCA AGT ACT GCG TAC-3’ and 5’-AAG GGG TTA TGC TAA AGC TTG CAT GCG GTA CTC AAT GGG CAC CTC GCG CGC GGG CAA TCT-3’ primers. The optimized RBS is underlined. The resulting fragment was then inserted by Gibson isothermal assembly into a KpnI-digested expression vector pKHT103 to generate pCF1193.

**pCF1328** [*aacC1 bla Tn7 lacI^q^ P_lac_::mltG E217Q*]. The plasmid pCF1328 for integration of an IPTG-inducible version of *mltG(E217Q)* catalytic mutant at the Tn7 attachment (*att*Tn7) site was created in several steps. First, the 5’ end of *mltG* gene was PCR-amplified from PAO1 gDNA with 5’-AGC TTA GTC GAC AGC TAG CCG GAT CCC CGG <u>GAG GAG GAT ACA T</u>AT GCG CAA ACT GCT GGT GCT GCT GGA GAG-3’ and *mltG E217Q* #2 (5’-TTC CGG CAC GCC GGT <u>TTG</u> CTT TTC CAC CAG CG-3’) primers. Concurrently, the 3’ end of *mltG* gene was PCR-amplified from PAO1 gDNA with *mltG E217Q* #1 (5’-CGC TGG TGG AAA AG<u>C AA</u>A CCG GCG TGC CGG AA-3’) and 5’-AAG GGG TTA TGC TAA AGC TTG CAT GCG GTA CTC ATT GTG GCG GCG GGG TGA TGG GC-3’ primers. The optimized RBS and the mutated site are underlined. These two PCR products were then combined by sewing PCR with 5’-AGC TTA GTC GAC AGC TAG CCG GAT CCC CGG <u>GAG GAG GAT ACA T</u>AT GCG CAA ACT GCT GGT GCT GCT GGA GAG-3’ and 5’-AAG GGG TTA TGC TAA AGC TTG CAT GCG GTA CTC ATT GTG GCG GCG GGG TGA TGG GC-3’ primers. This final PCR product was then inserted by Gibson isothermal assembly into a KpnI-digested expression vector pKHT103 to generate pCF1328.

For **pCF1009** [*aacC1 araC* P_ara_:: RBS_optimized_-*sltB1-FLAG*] and **pCF1010** [P_ara_:: RBS_optimized_-*sltB1 E135A-FLAG*], the *sltB1* genes were amplified from PAO1 gDNA or the appropriate mutated versions (pCF1193) using 5’-AAT TCC TGC AGC CCG GGG GAT CCA CTA GTT <u>GAG GAG</u> <u>GAT ACA T</u>GT GAA GAA CGC AAT GCA AGT ACT GCG TAC-3’ and 5’-TTG GAG CTC CAC CGC GGT GGC GGC CGC TCT AG**T CAC TTA TCA TCA TCA TCC TTA TAG TC**A TGG GCA CCT CGC GCG CGG GCA ATC T-3’ primers. The optimized RBS is underlined and the FLAG tag sequence is bolded. The resulting fragments were then inserted by Gibson isothermal assembly into a XbaI-digested expression vector pJN105 (7).

### *P. aeruginosa* strain construction

Briefly, during *P. aeruginosa* strain construction, plasmids were transferred into *P. aeruginosa* by conjugation from an *E. coli* donor [SM10(λpir)] on LB plates. Counter-selection against *E. coli* was accomplished on Vogel-Bonner minimal medium (VBMM)(11) supplemented with 30 μg/ml gentamicin.

To create the Δ*sltB1* strains CF1105 [PAO1Δ*sltB1*] and CF1143 [PAO1Δ*sltB1 attB::*P*_ampC_-lacZ*], pCF1097 [*aacC1 sacB oriT ‘PA4001-sltB1*Δ*(193-338)’*] was conjugated into PAO1 [WT] and CF263 [PAO1 *attB::*P*_ampC_-lacZ*] recipient from SM10(λpir) donor. For this purpose, PAO1 and CF263 were patched on an LB plate and grown overnight at 42°C while SM10(λpir) carrying pCF1097 was similarly grown at 37°C. Both the donor and the recipients were scraped, patched together onto an LB plate, and incubated at 37°C for ∼5h. The cells were scraped, resuspended in 500 μL of VBMM, diluted 1:10, and 100 μL of the resulting suspension was plated on VBMM supplemented with 30 μg/mL gentamicin. Plates were incubated at 37°C overnight. The exconjugants were purified on LB supplemented with 30 μg/mL gentamicin. A few single colonies were allowed to grow for ∼6h in plain LB broth to allow for the second plasmid recombination event, and 100 μL of the resulting culture was plated on LB supplemented with 5% (w/v) sucrose to select for the loss of the plasmid-encoded *sacB* gene. Sucrose-resistant colonies were then patched onto LB plates either containing or lacking 30 μg/mL gentamicin. Gentamicin-sensitive colonies were further tested by PCR with *sltB1*-flanking primers 5’-AAA TGT AAA GCA AGC TTC TGC AGG TCG ACT CTA GTT CAT GAA GCT GCT GAT GCC GAT GA-3’ and 5’-GAA TTC GAG CTC GAG CCC GGG GAT CCT TTT TTT TTT GAG TCG ATC TGC ACC GGC AGC-3’ to confirm gene deletion. The deletion retains the first one hundred ninety-two and last two codons of the *sltB1* reading frame. This truncation is similar to the one described in the previous study by Cavallari *et al*., which carries a SCAR mutation at nucleotide 577 of *sltB1* (12).

To create the Δ*slt* strains CF918 [PAO1Δ*slt*] and CF378 [PAO1Δ*sltB1*Δ*slt*], *slt* was deleted from PAO1 [WT] and CF1105 [PAO1Δ*sltB1*] by integration and re-circularization of pCF856 [*aacC1 sacB oriT ‘PA3020-slt*Δ*(3-635)’*] as described above. Sucrose-resistant, gentamicin-sensitive colonies were screened by PCR with *slt*-flanking primers 5’-AAC AGG CTG GAC TTG CCG GTA CCG TT-3’ and 5’-TTT TCT GGC CGT CGT TCA GGT GCT-3’. The deletion retains the first two and last eight codons of the *slt* reading frame.

To create the Δ*mltG* strains CF1410 [PAO1Δ*mltG*] and CF1416 [PAO1Δ*sltB1*Δ*mltG*] and CF1422 [PAO1Δ*sltB1*Δ*mltG* attB::P*_ampC_-lacZ*], *mltG* was deleted from PAO1 [WT], CF1105 [PAO1Δ*sltB1*] and CF1143 [PAO1Δ*sltB1 attB::*P*_ampC_-lacZ*] by integration and re-circularization of pCF1258 plasmid [*aacC1 sacB oriT* ‘*PA2963*-*mltG*Δ*(45-207)’*] as described above. The sucrose-resistant, gentamicin-sensitive colonies were screened by PCR with *mltG*-flanking primers 5’-AAA TGT AAA GCA AGC TTC TGC AGG TCG ACT CTA CTG GCC TAT GGC GAC GGG CTG TTC GAG A-3’ and 5’-GAA TTC GAG CTC GAG CCC GGG GAT CCT AAT GCT CGG CCA GGG CAC GCT TAC CGA T-3’. The deletion retains the first fourty-four and last one hundred fourty-three codons of the *mltG* reading frame.

To create the Δ*ampR* strains CF370 [PAO1Δ*sltB1*Δ*ampR*] and CF358 [PAO1Δ*sltB1*Δ*ampR attB::*P*_ampC_-lacZ*], *ampR* was deleted from CF1105 [PAO1Δ*sltB1*] and CF1143 [PAO1Δ*sltB1 attB::*P*_ampC_-lacZ*] by integration and re-circularization of pCF583 plasmid [*aacC1 sacB oriT* ‘*PA4109 ampR*Δ*(6-294)’*] as described above (2). Sucrose-resistant, gentamicin-sensitive colonies were screened by PCR with *ampR*-flanking primers 5’-AAC ACT TGC TGC TCC ATG AGC CGT TCG AA −3’ and 5’-AAG GTA TTC TTC TCG GCC CGC TCG AAG GT −3’. The deletion retains the first five and the last three codons of the *ampR* reading frame.

To create the Δ*ampG* strain CF372 [PAO1Δ*sltB1*Δ*ampG*], *ampG* was deleted from CF1105 [PAO1Δ*sltB1*] by integration and re-circularization of pCF284 plasmid [*aacC1 sacB oriT ‘PA4393-ampG*Δ*(1-594)’*] as described above (2). Sucrose-resistant, gentamicin-sensitive colonies were screened by PCR with *ampG*-flanking primers 5’-TAG AGC GGT TAG AGT GCG CGT TA-3’ and 5’-GTG CGA TCC ACG AAA AAG GC-3’. The deletion does not retain any *ampG* sequence.

To create the Δ*ampC* strain CF368 [PAO1Δ*sltB1*Δ*ampC*], *ampC* was deleted from CF1105 [PAO1Δ*sltB1*] by integration and re-circularization of pCF579 plasmid [*aacC1 sacB oriT ‘PA4110-ampC*Δ*(3-289)’*] as described above (2). Sucrose-resistant, gentamicin-sensitive colonies were screened by PCR with *ampC*-flanking primers 5’-ATG TCG ACG CGG TTG TTG TGG GTG GAC A −3’ and 5’-ATG GAA ATC CTC GCC GGC ATC CGC CTC −3’. The deletion retains the first three and the last nine codons of *ampC*.

Construction of strains CF666 [PAO1 *att*Tn7::*P_lac_::sltB1*] and CF1124 [PAO1 Δ*sltB1 att*Tn7::*P_lac_::sltB1*] with a P_lac_-regulated copy of *sltB1* (P*_lac_*_::_*sltB1*) integrated at Tn7 locus was based on a previously described protocol (11). In brief, plasmid pCF533, which encodes a P*_lac_*-regulated copy of *sltB1* flanked by Tn7 transposon inverted repeats, and plasmid pTNS2, which encodes Tn7 transposase, were co-electroporated into PAO1 [WT] and CF1105 [PAO1Δ*sltB1*]. Transformants were selected on LB plates supplemented with 30 μg/mL gentamicin. The integration of the transposon at the Tn7 attachment locus was confirmed by diagnostic PCR with PTn7R and PglmS-down primers (11). The gentamicin resistance cassette was then removed by Flp-mediated excision. Plasmid pFLP2 was electroporated and transformants were selected for growth on LB medium supplemented with 200 μg/mL carbenicillin, as described previously (11). Carbenicillin-resistant transformants were patched onto plain LB agar or LB supplemented with gentamicin to confirm the loss of the gentamicin resistance cassette. Gentamicin sensitive clones were grown overnight in liquid LB medium lacking antibiotics and purified on LB agar supplemented with 5% (w/v) sucrose to select for the loss of the pFLP2 plasmid, which encodes the *sacB* gene. Isolated colonies were patched onto LB supplemented with carbenicillin or no antibiotic to confirm the loss of the pFLP2 plasmid. The construction of strain CF1122 [PAO1 Δ*sltB1 att*Tn7::*P_lac_::empty*] and CF1212 [PAO1 Δ*sltB1 att*Tn7::*P_lac_::sltB1 E135A*], with P*_lac_*::empty and P*_lac_*::*sltB1* integrated at the Tn7 locus were performed as above in strain CF1105 [PAO1Δ*sltB1*], but using plasmids pKHT103 and pCF1193, respectively.

Construction of strains CF1447 [PAO1 Δ*sltB1*Δ*mltG att*Tn7::*P_lac_::empty*], CF1449 [PAO1 Δ*sltB1*Δ*mltG att*Tn7::*P_lac_::mltG*] and CF1451 [PAO1 Δ*sltB1*Δ*mltG att*Tn7::*P_lac_::mltG E217Q*] with with P*_lac_*::empty, P*_lac_*::*mltG* and P*_lac_*::*mltG E217Q* integrated at the Tn7 locus were performed as above in strain CF1416 [PAO1Δ*sltB1*ΔmltG] using plasmids pKHT103, pCF658 and pCF1328, respectively.

Construction of strains CF1304 [PAO1 Δ*sltB1*Δ*mltG attB::*P*_ampC_-lacZ att*Tn7::*P_lac_::empty*], CF1306 [PAO1 Δ*sltB1*Δ*mltG attB::*P*_ampC_-lacZ att*Tn7::*P_lac_::mltG*] and CF1308 [PAO1 Δ*sltB1*Δ*mltG attB::*P*_ampC_-lacZ att*Tn7::*P_lac_::mltG E217Q*] with P*_lac_*::empty, P*_lac_*::*mltG* and P*_lac_*::*mltG E217Q* integrated at the Tn7 locus were performed as above in strain CF1422 [PAO1Δ*sltB1*Δ*mltG attB::*P*_ampC_-lacZ*] using plasmids pKHT103, pCF658 and pCF1328, respectively.

Strains CF1631 [PAO1 (empty)], CF1632 [PAO1 (P_ara_-*sltB1*-FLAG)] and CF1634 [PAO1 (P_ara_-*sltB1 E135A*-FLAG)] were obtained by electroporation of plasmids pJN105 (7), pCF1009 and pCF1010 in PAO1 [WT], respectively.

Strains CF1726 [Δ*ampC* (empty)], CF1727 [Δ*ampC* (P_ara_-*sltB1*-FLAG)] and CF1728 [Δ*ampC* (P_ara_-*sltB1 E135A*-FLAG)] were obtained by electroporation of plasmids pJN105 (7), pCF1009 and pCF1010 in CF612 [Δ*ampC*] (2), respectively.

Strains CF1731 [Δ*ampR* (empty)], CF1732 [Δ*ampR* (P_ara_-*sltB1*-FLAG)] and CF1733 [Δ*ampR* (P_ara_-*sltB1 E135A*-FLAG)] were obtained by electroporation of plasmids pJN105 (7), pCF1009 and pCF1010 in CF550 [Δ*ampR*] (2), respectively.

Strains CF1738 [Δ*mltG* (empty)], CF1739 [Δ*mltG* (P_ara_-*sltB1*-FLAG)] and CF1740 [Δ*mltG* (P_ara_-*sltB1 E135A*-FLAG)] were obtained by electroporation of plasmids pJN105 (7), pCF1009 and pCF1010 in CF1410 [Δ*mltG*], respectively.

## REFERENCES

1. Strateva T, Yordanov D. 2009. Pseudomonas aeruginosa – a phenomenon of bacterial resistance. Journal of Medical Microbiology.58:1133–1148.

2. López-Causapé C, Cabot G, Del Barrio-Tofiño E, Oliver A. 2018. The Versatile Mutational Resistome of Pseudomonas aeruginosa. Front Microbiol 9:685.

3. Fisher JF, Mobashery S. 2014. The sentinel role of peptidoglycan recycling in the β-lactam resistance of the Gram-negative Enterobacteriaceae and Pseudomonas aeruginosa. Bioorganic Chemistry 56:41–48.

4. Livermore DM. 1995. Beta-Lactamases in laboratory and clinical resistance. Clinical Microbiology Reviews 8:557–584.

5. Livermore DM. 1987. Clinical significance of beta-lactamase induction and stable derepression in gram-negative rods. Eur J Clin Microbiol 6:439–445.

6. Giwercman B, Lambert PA, Rosdahl VT, Shand GH, Heiby N. 1990. Rapid emergence of resistance in pseudomonas aeruginosa in cystic fibrosis patients due to in-vivo selection of stable partially derepressed β-lactamase producing strains. Journal of Antimicrobial Chemotherapy 26:247–259.

7. Juan C, Maciá MD, Gutiérrez O, Vidal C, Pérez JL, Oliver A. 2005. Molecular mechanisms of β-lactam resistance mediated by AmpC hyperproduction in Pseudomonas aeruginosa clinical strains. Antimicrobial Agents and Chemotherapy 49:4733–4738.

8. Moya B, Dötsch A, Juan C, Blázquez J, Zamorano L, Haussler S, Oliver A. 2009. β-lactam resistance response triggered by inactivation of a nonessential penicillin-binding protein. PLoS Pathogens 5:e1000353.

9. Moyá B, Beceiro A, Cabot G, Juan C, Zamorano L, Alberti S, Oliver A. 2012. Pan-β-lactam resistance development in Pseudomonas aeruginosa clinical strains: Molecular mechanisms, penicillin-binding protein profiles, and binding affinities. Antimicrobial Agents and Chemotherapy 56:4771–4778.

10. Jacobs C, Huang LJ, Bartowsky E, Normark S, Park JT. 1994. Bacterial cell wall recycling provides cytosolic muropeptides as effectors for beta-lactamase induction. The EMBO journal 13:4684–4694.

11. Boudreau MA, Fisher JF, Mobashery S. 2012. Messenger functions of the bacterial cell wall-derived muropeptides. Biochemistry 51:2974–2990.

12. Lee M, Dhar S, Debenedetti S, Hesek D, Boggess B, Blázquez B, Mathee K, Mobashery S. 2016. Muropeptides in Pseudomonas aeruginosa and their Role as Elicitors of β-Lactam-Antibiotic Resistance. Angewandte Chemie 55:6882–6886.

13. Dik DA, Domínguez-Gil T, Lee M, Hesek D, Byun B, Fishovitz J, Boggess B, Hellman LM, Fisher JF, Hermoso JA, Mobashery S. 2017. Muropeptide Binding and the X-ray Structure of the Effector Domain of the Transcriptional Regulator AmpR of Pseudomonas aeruginosa. J Am Chem Soc 139:1448–1451.

14. Rohs PDA, Bernhardt TG. 2021. Growth and Division of the Peptidoglycan Matrix. Annu Rev Microbiol 75:315–336.

15. Cho H, Wivagg CN, Kapoor M, Barry Z, Rohs PDA, Suh H, Marto JA, Garner EC, Bernhardt TG. 2016. Bacterial cell wall biogenesis is mediated by SEDS and PBP polymerase families functioning semi-autonomously. Nat Microbiol 1:16172.

16. Meeske AJ, Riley EP, Robins WP, Uehara T, Mekalanos JJ, Kahne D, Walker S, Kruse AC, Bernhardt TG, Rudner DZ. 2016. SEDS proteins are a widespread family of bacterial cell wall polymerases. Nature 537:634–638.

17. Rohs PDA, Buss J, Sim SI, Squyres GR, Srisuknimit V, Smith M, Cho H, Sjodt M, Kruse AC, Garner EC, Walker S, Kahne DE, Bernhardt TG. 2018. A central role for PBP2 in the activation of peptidoglycan polymerization by the bacterial cell elongation machinery. PLoS Genet 14:e1007726.

18. Dörr T, Cava F, Lam H, Davis BM, Waldor MK. 2013. Substrate specificity of an elongation-specific peptidoglycan endopeptidase and its implications for cell wall architecture and growth of Vibrio cholerae. Molecular Microbiology 89:949–962.

19. Singh SK, Saisree L, Amrutha RN, Reddy M. 2012. Three redundant murein endopeptidases catalyse an essential cleavage step in peptidoglycan synthesis of Escherichia coli K12. Molecular Microbiology 86:1036–1051.

20. Hashimoto M, Ooiwa S, Sekiguchi J. 2012. Synthetic lethality of the lytE cwlO genotype in Bacillus subtilis is caused by lack of D, L-endopeptidase activity at the lateral cell wall. Journal of Bacteriology 194:796–803.

21. Bisicchia P, Noone D, Lioliou E, Howell A, Quigley S, Jensen T, Jarmer H, Devine KM. 2007. The essential YycFG two-component system controls cell wall metabolism in Bacillus subtilis. Mol Microbiol 65:180–200.

22. Tipper DJ, Strominger JL. 1965. Mechanism of action of penicillins: a proposal based on their structural similarity to acyl-D-alanyl-D-alanine. Proceedings of the National Academy of Sciences 54:1133–1141.

23. Cho H, Uehara T, Bernhardt TG. 2014. Beta-lactam antibiotics induce a lethal malfunctioning of the bacterial cell wall synthesis machinery. Cell 159:1300–1311.

24. Dik DA, Marous DR, Fisher JF, Mobashery S. 2017. Lytic transglycosylases: concinnity in concision of the bacterial cell wall. Critical Reviews in Biochemistry and Molecular Biology 52:503–542.

25. Goodell EW. 1985. Recycling of Murein by Escherichia coli. Journal of Bacteriology 163:305–310.

26. Park JT, Uehara T. 2008. How Bacteria Consume Their Own Exoskeletons (Turnover and Recycling of Cell Wall Peptidoglycan). Microbiology and Molecular Biology Reviews 72:211–227.

27. Caille O, Zincke D, Merighi M, Balasubramanian D, Kumari H, Kong KF, Silva-Herzog E, Narasimhan G, Schneper L, Lory S, Mathee K. 2014. Structural and functional characterization of Pseudomonas aeruginosa global regulator AmpR. Journal of Bacteriology 196:3890–3902.

28. Fumeaux C, Bernhardt TG. 2017. Identification of MupP as a New Peptidoglycan Recycling Factor and Antibiotic Resistance Determinant in Pseudomonas aeruginosa. mBio 8.

29. Yunck R, Cho H, Bernhardt TG. 2016. Identification of MltG as a potential terminase for peptidoglycan polymerization in bacteria. Mol Microbiol 99:700–718.

30. Sonnabend MS, Klein K, Beier S, Angelov A, Kluj R, Mayer C, Groß C, Hofmeister K, Beuttner A, Willmann M, Peter S, Oberhettinger P, Schmidt A, Autenrieth IB, Schütz M, Bohn E. 2020. Identification of drug resistance determinants in a clinical isolate of pseudomonas aeruginosa by high-density transposon mutagenesis. Antimicrobial Agents and Chemotherapy 64:e01771–19.

31. Dik DA, Madukoma CS, Tomoshige S, Kim C, Lastochkin E, Boggess WC, Fisher JF, Shrout JD, Mobashery S. 2019. Slt, MltD, and MltG of Pseudomonas aeruginosa as Targets of Bulgecin A in Potentiation of β-Lactam Antibiotics. ACS Chem Biol 14:296– 303.

32. Cavallari JF, Lamers RP, Scheurwater EM, Matos AL, Burrows LL. 2013. Changes to its peptidoglycan-remodeling enzyme repertoire modulate β-lactam resistance in Pseudomonas aeruginosa. Antimicrobial Agents and Chemotherapy 57:3078–3084.

33. Lamers RP, Nguyen UT, Nguyen Y, Buensuceso RNC, Burrows LL. 2015. Loss of membrane-bound lytic transglycosylases increases outer membrane permeability and β-lactam sensitivity in Pseudomonas aeruginosa. MicrobiologyOpen 4:879–895.

34. Kong K-F, Jayawardena SR, Indulkar SD, Puerto AD, Koh C-L, Høiby N, Mathee K. 2005. Pseudomonas aeruginosa AmpR is a global transcriptional factor that regulates expression of AmpC and PoxB beta-lactamases, proteases, quorum sensing, and other virulence factors. Antimicrobial Agents and Chemotherapy 49:4567–4575.

35. Jorgenson MA, Chen Y, Yahashiri A, Popham DL, Weiss DS. 2014. The bacterial septal ring protein RlpA is a lytic transglycosylase that contributes to rod shape and daughter cell separation in Pseudomonas aeruginosa. Mol Microbiol 93:113–128.

36. Kong KF, Aguila A, Schneper L, Mathee K. 2010. Pseudomonas aeruginosa β-lactamase induction requires two permeases, AmpG and AmpP. BMC microbiology 10:328.

37. Zamorano L, Reeve TM, Juan C, Moya B, Cabot G, Vocadlo DJ, Mark BL, Oliver A. 2011. AmpG Inactivation Restores Susceptibility of Pan- -Lactam-Resistant Pseudomonas aeruginosa Clinical Strains. Antimicrobial Agents and Chemotherapy 55:1990–1996.

38. Holtje JV. 1998. Growth of the stress-bearing and shape-maintaining murein sacculus of Escherichia coli. Microbiology and Molecular Biology Reviews : MMBR 62:181–203.

39. Legaree BA, Clarke AJ. 2008. Interaction of Penicillin-Binding Protein 2 with Soluble Lytic Transglycosylase B1 in Pseudomonas aeruginosa. Journal of Bacteriology 190:6922–6926.

40. Nikolaidis I, Izoré T, Job V, Thielens N, Breukink E, Dessen AA, Izore T, Job V, Thielens N, Breukink E, Dessen AA. 2012. Calcium-dependent complex formation between PBP2 and lytic transglycosylase SltB1 of Pseudomonas aeruginosa. Microbial Drug Resistance 18:298–305.

41. Huang Y-W, Wu C-J, Hu R-M, Lin Y-T, Yang T-C. 2015. Interplay among Membrane-Bound Lytic Transglycosylase D1, the CreBC Two-Component Regulatory System, the AmpNG-AmpDI-NagZ-AmpR Regulatory Circuit, and L1/L2 β-Lactamase Expression in Stenotrophomonas maltophilia. Antimicrob Agents Chemother 59:6866–6872.

42. Yin J, Sun Y, Sun Y, Yu Z, Qiu J, Gao H. 2018. Deletion of Lytic Transglycosylases Increases Beta-Lactam Resistance in Shewanella oneidensis. Frontiers in Microbiology 9:13.

43. Choi K-H, Kumar A, Schweizer HP. 2006. A 10-min method for preparation of highly electrocompetent Pseudomonas aeruginosa cells: application for DNA fragment transfer between chromosomes and plasmid transformation. J Microbiol Methods 64:391–397.

44. Jacobs MA, Alwood A, Thaipisuttikul I, Spencer D, Haugen E, Ernst S, Will O, Kaul R, Raymond C, Levy R, Chun-Rong L, Guenthner D, Bovee D, Olson MV, Manoil C. 2003. Comprehensive transposon mutant library of Pseudomonas aeruginosa. Proceedings of the National Academy of Sciences 100:14339–14344.

45. Altschul SF, Gish W, Miller W, Myers EW, Lipman DJ. 1990. Basic local alignment search tool. J Mol Biol 215:403–410.

46. Stover CK, Pham XQ, Erwin AL, Mizoguchi SD, Warrener P, Hickey MJ, Brinkman FS, Hufnagle WO, Kowalik DJ, Lagrou M, Garber RL, Goltry L, Tolentino E, Westbrock-Wadman S, Yuan Y, Brody LL, Coulter SN, Folger KR, Kas A, Larbig K, Lim R, Smith K, Spencer D, Wong GK, Wu Z, Paulsen IT, Reizer J, Saier MH, Hancock RE, Lory S, Olson MV. 2000. Complete genome sequence of Pseudomonas aeruginosa PAO1, an opportunistic pathogen. Nature 406:959–964.

47. Bradford MM. 1976. A rapid and sensitive method for the quantitation of microgram quantities of protein utilizing the principle of protein-dye binding. Anal Biochem 72:248– 254.

48. Vadlamani G, Thomas MD, Patel TR, Donald LJ, Reeve TM, Stetefeld J, Standing KG, Vocadlo DJ, Mark BL. 2015. The β-lactamase gene regulator AmpR is a tetramer that recognizes and binds the D-Ala-D-Ala motif of its repressor UDP-N-acetylmuramic acid (MurNAc)-pentapeptide. Journal of Biological Chemistry 290:2630–2643.

49. Ropy A, Cabot G, Sánchez-Diener I, Aguilera C, Moya B, Ayala JA, Oliver A. 2015. Role of Pseudomonas aeruginosa low-molecular-mass penicillin-binding proteins in AmpC expression, β-lactam resistance, and peptidoglycan structure. Antimicrobial Agents and Chemotherapy 59:3925–3934.

## References for supplemental materials

1. Stover CK, Pham XQ, Erwin AL, Mizoguchi SD, Warrener P, Hickey MJ, Brinkman FS, Hufnagle WO, Kowalik DJ, Lagrou M, Garber RL, Goltry L, Tolentino E, Westbrock-Wadman S, Yuan Y, Brody LL, Coulter SN, Folger KR, Kas A, Larbig K, Lim R, Smith K, Spencer D, Wong GK, Wu Z, Paulsen IT, Reizer J, Saier MH, Hancock RE, Lory S, Olson MV. 2000. Complete genome sequence of Pseudomonas aeruginosa PAO1, an opportunistic pathogen. Nature 406:959–964.

2. Fumeaux C, Bernhardt TG. 2017. Identification of MupP as a New Peptidoglycan Recycling Factor and Antibiotic Resistance Determinant in Pseudomonas aeruginosa. mBio 8.

3. Simon R, Priefer U, Pühler A. 1983. A Broad Host Range Mobilization System for In Vivo Genetic Engineering: Transposon Mutagenesis in Gram Negative Bacteria. Bio/Technology 1:784–791.

4. Rietsch A, Vallet-Gely I, Dove SL, Mekalanos JJ. 2005. ExsE, a secreted regulator of type III secretion genes in Pseudomonas aeruginosa. Proc Natl Acad Sci U S A 102:8006–8011.

5. Hoang TT, Karkhoff-Schweizer RR, Kutchma AJ, Schweizer HP. 1998. A broad-host-range Flp-FRT recombination system for site-specific excision of chromosomally-located DNA sequences: application for isolation of unmarked Pseudomonas aeruginosa mutants. Gene 212:77–86.

6. Choi K-H, Gaynor JB, White KG, Lopez C, Bosio CM, Karkhoff-Schweizer RR, Schweizer HP. 2005. A Tn7-based broad-range bacterial cloning and expression system. Nat Methods 2:443–448.

7. Newman JR, Fuqua C. 1999. Broad-host-range expression vectors that carry the L-arabinose-inducible Escherichia coli araBAD promoter and the araC regulator. Gene 227:197–203.

8. Vieira J, Messing J. 1991. New pUC-derived cloning vectors with different selectable markers and DNA replication origins. Gene 100:189–194.

9. Caille O, Zincke D, Merighi M, Balasubramanian D, Kumari H, Kong KF, Silva-Herzog E, Narasimhan G, Schneper L, Lory S, Mathee K. 2014. Structural and functional characterization of Pseudomonas aeruginosa global regulator AmpR. Journal of Bacteriology 196:3890–3902.

10. Gibson DG, Young L, Chuang R-Y, Venter JC, Hutchison CA 3rd, Smith HO. 2009. Enzymatic assembly of DNA molecules up to several hundred kilobases. Nat Methods 6:343–345.

11. Choi K-H, Schweizer HP. 2006. mini-Tn7 insertion in bacteria with single attTn7 sites: example Pseudomonas aeruginosa. Nat Protoc 1:153–161.

12. Cavallari JF, Lamers RP, Scheurwater EM, Matos AL, Burrows LL. 2013. Changes to its peptidoglycan-remodeling enzyme repertoire modulate β-lactam resistance in Pseudomonas aeruginosa. Antimicrobial Agents and Chemotherapy 57:3078–3084.

